# Machine Learning Reveals Modules of Economic Behavior from Foraging Mice

**DOI:** 10.1101/357434

**Authors:** Cornelia N Stacher Hörndli, Eleanor Wong, Elliott Ferris, Alexis Nikole Rhodes, P. Thomas Fletcher, Christopher Gregg

**Author notes:** Correspondence: Christopher Gregg, PhD. Assistant Professor, Neurobiology and Anatomy and Human Genetics Room 408B, Biopolymers Research Building Bld. 570 20 South 2030 East University of Utah Salt Lake City, UT 84112.

## Abstract

Complex ethological behaviors could be constructed from modules that are discrete functional units of behavior with a genetic basis. Here, we test this idea for foraging, and develop a paradigm to dissect foraging patterns in mice. We uncover discrete behavioral modules linked to round trip excursions from the home. Machine learning reveals 59 modules across different genetic backgrounds and ages. Different modules develop at different ages and are linked to different aspects of economic behavior, including memory, reward, risk and effort responses. Crosses of distant mouse strains reveal that parental and genetic effects shape foraging differently, and parental effects grow stronger with age. Specific behavioral modules, genes and pathways are found to be sensitive to parental effects. One candidate gene, *Magel2*, is linked to Prader-Willi Syndrome and shaped the expression of discrete modules in an age-dependent manner. Our results reveal building blocks for normal and abnormal economic behavior patterns.

**HIGHLIGHTS:**

- Identification of 59 economic behavior modules underlying foraging
- Discrete modules are linked to memory, reward, risk and effort responses
- Genetic and parental effects shape foraging by changing module expression
- *Magel2*, a Prader-Willi Syndrome gene, affects specific modules at specific ages

## INTRODUCTION

Most genomic elements influencing mammalian brain development and function shape instinctive and learned behavior patterns in ways that are unknown (Bakken et al., 2016; Kang et al., 2011; Miller et al., 2014; Silbereis et al., 2016; Thompson et al., 2014). Behavior could be constructed from discrete “modules” that are independent units of behavior (Vogelstein et al., 2014; Wiltschko et al., 2015; Yang and Shah, 2014). From this perspective, specific genomic elements could have evolved to shape specific behavioral modules, which in turn are building blocks for more complex patterns. Changes to the genome, and brain structure and function, would thereby shape new behavior patterns by either altering the expression of existing modules and/or by creating novel modules. However, for most complex ethological behaviors, methods are lacking to test this hypothesis. It is generally unclear how to define modules. We do not know which behavioral patterns are comprised of modules, the identity of the modules involved, how they are expressed or the mechanisms involved. Module expression potentially encompasses frequency, timing and sequential order. There are three main problems to solve. First, improved methods are needed to study complex, ethological behavior patterns. Second, methods are needed that dissect apart these patterns to identify underlying modules. Finally, approaches are needed to investigate how module structure, module expression and behavior patterns change in response to targeted genetic/environmental manipulations.

Here, we devise methods to investigate potential modular architecture in foraging, a rich and deeply conserved repertoire of economic behavioral responses (Stephens et al., 2007). Many neural systems are involved in foraging, including novelty-seeking, anxiety, reward, perseveration, sensorimotor integration, hunger, satiety, attention, activity, navigation, as well as learning and memory (Stephens et al., 2007). Natural selection shapes foraging patterns through these systems to minimize predation risk and energy expenditure, while maximizing caloric intake (Brown and Laundré, 1999; Pulliam and Charnov, 1977; Stephens et al., 2007). Understanding the basis of different foraging patterns and tradeoffs is considered important for understanding mammalian economic behaviors (Brown and Laundré, 1999; Pulliam and Charnov, 1977; Stephens et al., 2007), which have roles in human obesity, addiction, fear, anxiety and psychiatric disorders (Monterosso et al., 2012; Rowland et al., 2008; Sharp et al., 2012a).

Learning how complex foraging patterns are constructed at different ages could also advance our understanding of the principles of behavioral development. Selective pressures on foraging shaped mammalian life histories, including brain and behavioral developmental processes (Barton, 2012; Jones, 2011; Melin et al., 2014; Reader and Laland, 2002; Sol et al., 2005; Walker et al., 2006). Indeed, major mammalian developmental milestones are linked to foraging, such as the elimination of suckling during nursing and emergence of independent foraging after weaning (Galef, 1981; Hewlett and Lamb, 2010).

Our study tests the hypothesis that foraging is comprised of discrete behavioral modules differentially expressed at different ages and in response to genetic and/or parental effects. We further test for the existence of distinct modules shaping memory, reward, risk and effort responses. To achieve this, we introduce a rich foraging behavior paradigm, machine-learning approaches to study foraging architecture in mice and an approach to detect modular architecture from round trip foraging excursions from a home base. Parental and genetic effects on foraging, module formation and module expression are investigated at different ages using reciprocal F1 hybrid offspring derived from crosses of distantly inbred subspecies of *Mus musculus*. The expression of specific modules changes due to parental and genetic effects in an age-dependent manner, and for parental effects, we perform genomics and functional studies to uncover genes involved. Top candidates are genes linked to Prader-Willi Syndrome (PWS) and autism (Cassidy et al., 2012; Schaaf et al., 2013). We perform functional validation studies for one of these genes, *Magel2*, to test for age-dependent effects on specific modules and gain deeper insights into how loss of *Magel2* function shapes behavior. Overall, our study uncovers an atlas of 59 core modules underlying mouse foraging patterns across different ages, helping to elucidate behavioral development, the basis of normal and abnormal economic behavior patterns and how evolution shapes new patterns.

## RESULTS

### An Approach to Uncover Modular Architecture in Mouse Foraging

Behavioral patterns encompass where, when and what an animal does, while behavioral modules are the building blocks for *how* it constructs the pattern. To test for foraging modules, we required a rich assay in which mice exhibit diverse, ethological foraging patterns and methods to identify underlying behavioral modules. It is challenging to recapitulate all ecological elements of foraging in the lab. One element frequently overlooked is the home base (Eilam and Golani, 1989). For animals in the wild, the home base is a refuge from predation, shelter from the environment, and a safe haven for rest, mating, offspring rearing and food storage. Selective pressures related to the home can shape behavior patterns (Eilam and Golani, 1989; Gorny et al., 2002; Weber et al., 2013). Most lab tests miss this dimension of behavior and lift the animal from the home cage into a testing apparatus (Wahlsten, 2010). Others monitor home cage behaviors, but in the absence of access to an outside environment (Loos et al., 2014). Therefore, existing assays are unsuitable for our objectives.

The central role of the home base in rodent and human ecology potentially makes it a key anchor point to discover modular structure in complex foraging patterns. In previous studies rodents performed variable types of round trip excursions when a home base was available during an open field test (Eilam and Golani, 1989). We hypothesized that a richer assay, designed to allow the expression of diverse and complex foraging patterns, would reveal a deeper variety of home base excursion subtypes. Our rationale was that subtypes of foraging excursions from the home base would be discrete modules of economic behavior, involving different motivations, functions and underlying mechanisms. The expression frequency, timing and sequence of these modules would then shape more complex foraging patterns over time. Alternatively, there may be near-infinite types of excursions composed of different behavioral responses in a random order.

Based on this rationale, we designed a foraging paradigm that examines a rich behavioral landscape, while permitting free access to a home base cage (Fig. 1A, see Supplemental Information). The paradigm integrates concepts from behavioral ecology, including sand and food patches containing seeds (Brown and Kotler, 1994; Stephens et al., 2007). We constructed foraging arenas with 4 pots filled with sand, one of which also contains seeds (the food patch) (Fig. 1A-C). For the home base, mouse cages are fitted with a 3D printed tunnel that is “plugged” into a connecting tunnel to the foraging arena (Fig. 1A). The experimenter starts a trial by removing a door in the tunnel, allowing access to the arena. This reduces handling and promotes innate, ethological behavior sequences. Behavior testing encompasses two phases: a novel Exploration phase and a familiar Foraging phase (Fig. 1B-D), as described below. A protocol to print, assemble and run the assay is provided (Supplemental Information).

**Figure 1.**
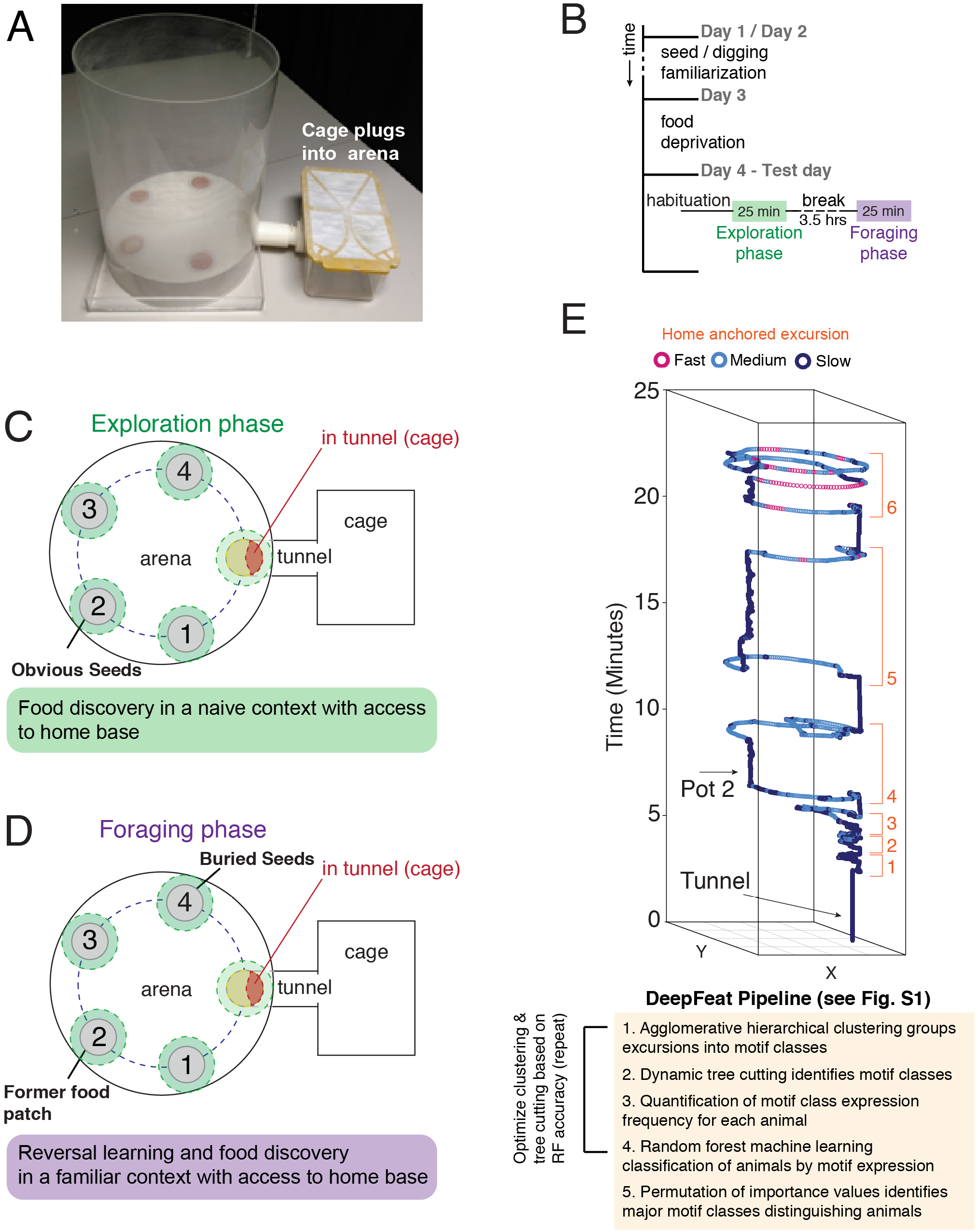
A paradigm to study rich foraging patterns in mice. **(A)** Image shows the foraging arena. The 3D printed arena platform holds pots containing sand and one food patch with seeds. The home base cage plugs into the arena through a 3D printed tunnel on the side. Access to the arena is controlled by a door in the tunnel and the experimenter. **(B)** A summary of the assay workflow. **(C and D)** Overviews of the pots, tunnel zone and seed location during the naïve Exploration phase (C) and familiar Foraging phase (D). The mice learn the food patch is Pot#2 in the Exploration phase. In the Foraging phase, the food is moved and hidden in Pot#4. **(E)** An example of XY tracking of body position over time during the foraging assay (Exploration phase shown). Excursions are defined from round trips from home base cage (orange numbered track sections). Foraging behavior modules are defined from these excursions using DeepFeat (see text, Fig. S1 and Supplemental Information).

During the Exploration phase, mice are given 25 minutes to spontaneously enter the arena, explore and discover seeds placed on top of the sand in the food patch at Pot#2 (Fig. 1B,C). During this phase, the mouse expresses behavioral sequences related to the exploration of a novel environment and the discovery and consumption of food in a novel food patch (Movie S1). Four hours later, the mice are again given 25 minutes of access to the arena in the Foraging Phase. However, the seeds are no longer atop Pot#2 but are buried in the sand in Pot#4 (Fig. 1B,D). During this more familiar and goal-directed phase, mice express behavioral sequences related to their expectation of the former food patch in the now familiar environment and the discovery of a new, hidden food source (Pot#4) (Movie S2). We food deprived the mice prior to the assay such that the animals experience the same degree of body weight loss (~10% of starting body weight), setting a common baseline for comparison across ages (see Supplemental Information and Experimental Procedures). Food deprivation increases learning and stereotyped patterns of behavior (Makowiecki et al., 2012) (Fig. 1B).

### Resolving Foraging Patterns in Mice

To analyze behavior patterns, we defined features that describe where, when and what an animal does during testing. The features systematically atomize foraging into the spatiotemporal components that show where the animals go in the arena during the Exploration and Foraging phases, quantifying their visits to different areas (Fig. 1-S1A,B). Spatiotemporal features are further broken into 5 minute time bins that capture patterns from five different early, mid and late stages temporal windows. Finally, we measure features germane to foraging, which include total food consumption, digging and distance traveled (Fig. 1-S1B). Locomotor features related to velocity, numbers of excursions performed and gait patterns (slow, medium, fast) are also measured. In total, foraging patterns are deconstructed into 338 features that are each assigned a unique ID (Fig. 1-S1B and Table S1 and S2).

We tested whether this feature space captures diverse and complex (high dimensional) foraging patterns in our paradigm. In the study detailed below, we collect foraging profiles for 383 mice of different ages, genotypes and crosses. Principal component analysis of the feature data from all 383 mice revealed high dimensionality (Fig. 1-S1D,E). A parallel analysis found that twenty-four principal components are retained to explain the variance in the data (Fig. 1-S1E). Therefore, features and assay reveal complex behavioral patterns, as intended. Below, we build on these foundations to apply supervised machine-learning and data visualization strategies to uncover foraging patterns for different ages, genotypes and crosses.

### Discovering the Modules Underlying Foraging Patterns

Our final major methodological challenge was to devise an approach to uncover modules that are discrete units of behavior underlying more complex foraging patterns. As described above, we hypothesize that modules could be defined as home base anchored excursions (Fig. 1E). To potentially identify these modules, we developed a data analysis pipeline in R, called “DeepFeat”, which takes XY tracking data from commercially available tracking software as an input. DeepFeat detects discrete modules from excursions relative to a user-defined anchor point (the home base) (Fig. 1E and Supplemental Information). For each excursion, DeepFeat extracts a vector of statistical features describing XY movements over time. We refer to these as “phenovectors”. For all excursions performed by a cohort of mice, agglomerative hierarchical clustering is performed on the phenovectors and a dynamic tree cutting algorithm (Langfelder et al., 2008) is used to define discrete clusters of related excursions in a dendrogram. A cut branch in the dendrogram thereby defines excursions of a similar type, which collectively comprise a discrete module of behavior. Individual excursions are assigned a module identification key that we call a Concise Idiosyncratic Module Alignment Report (CIMAR) string. For each excursion, the module assignment, underlying raw data and metadata (eg. mouse of origin, time of expression, phase and other details), are stored in a list with CIMAR string keys, allowing simple data retrieval, quantification and analysis of the expression of different modules.

To test whether particular phenovectors and clustering conditions identify bona fide, biologically relevant modules, we use random forest machine learning to determine whether mice in a cohort can be classified from the expression of the identified modules according to a factor of interest (eg. age, genotype, cross, etc). Therefore, similar to recursive methods for feature selection in the machine-learning field (Miao and Niu, 2016), our approach can be tuned to optimize module detection to maximize classification accuracy (Fig. 1-S2). The generalization error of the classifier is conservatively estimated using out-of-bag (OOB) error (Breiman, 1996). This process detects bona fide, biological modules of behavior, as shown below. We now turn to the application of all these methods to first study foraging patterns.

**Figure 2.**
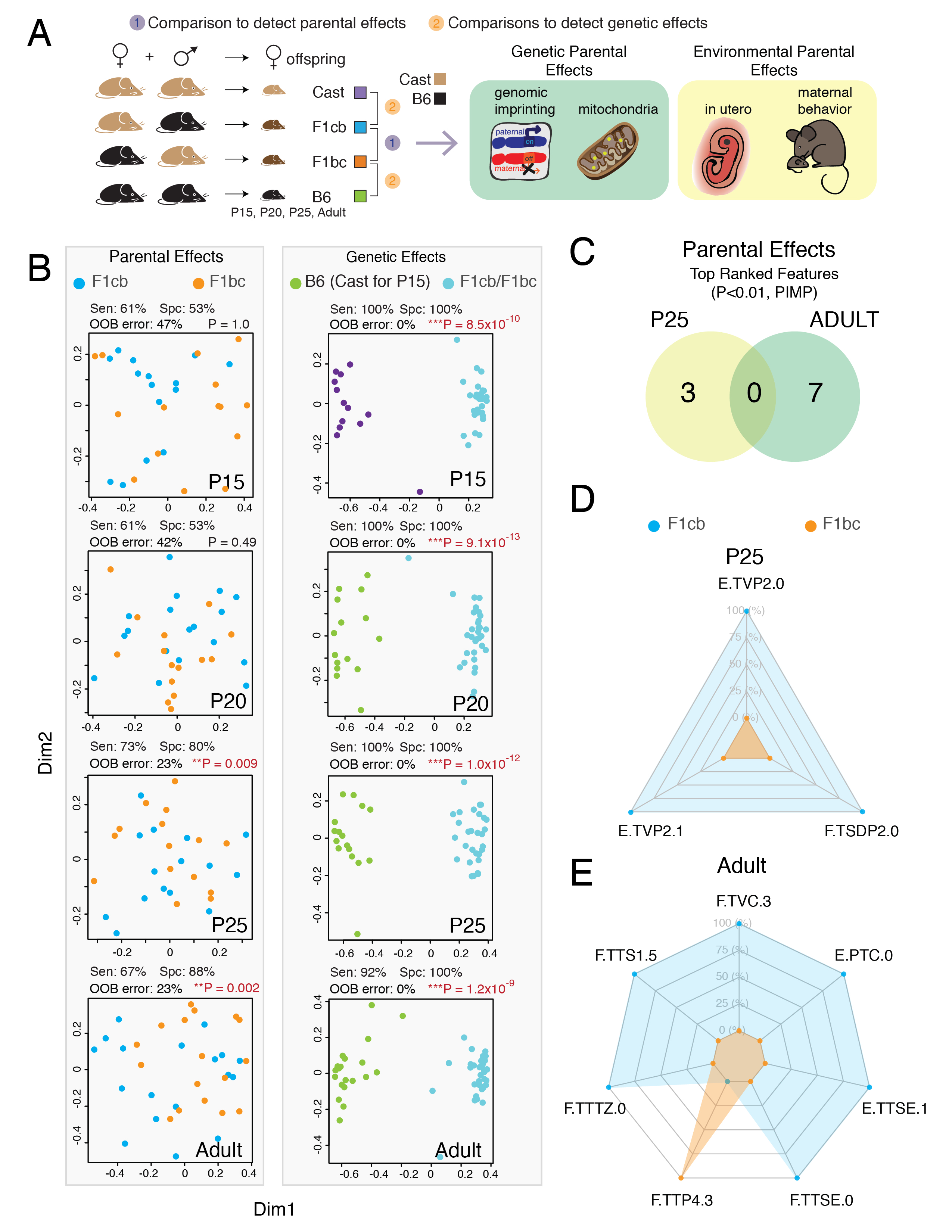
Random forest analysis of foraging patterns reveals parental effects on economic patterns emerge after weaning, while genetic effects are robust at all ages. (**A**) Schematic summary of mouse genetics strategy to identify parental and/or genetic effects at P15, P20, P25 and adulthood. Different mice are tested at each age. Parental effects potentially encompass genetic parental effects and environmental parental effects (see text). (**B**) Multidimensional scaling plots of the random forest classifier proximity values for each mouse reveals parental and/or genetic effects in P15 (early weaning), P20 (weaning), P25 (post-weaning) and adult females. The sensitivity (Sen), specificity (Spc), out-of-bag error (OOB) and two-tailed Fisher test P-value for the confusion matrix of the classification results are shown above each plot. (**C**) Venn diagram shows that the significantly informative features in the classifiers detecting parental effects in P25 and adult F1cb vs F1bc mice differ (PIMP P<0.01). (**D and E**) Radar plots show the data for significantly informative features for the classifiers detecting parental effects in P25 (D) and adult (E) offspring. The results show the relative values for the features that most strongly distinguish F1cb and F1bc offspring foraging patterns at each age (see main text). Feature data points are the mean values from F1cb (blue) and F1bc (orange) mice, presented as percentages of the relative maximum (100%) and minimum (0%) values for each feature. Feature codes are: [Phase.Feature.TimeBin]. F, Foraging phase; E, Exploration phase. See Table S1 for feature definitions.

### Parental Effects in Mice Shape Foraging Patterns at Later Ages, While Genetic Effects Are Robust from Weanling to Adulthood

We hypothesized that parental and genetic effects shape different foraging patterns in offspring. Independent foraging emerges at weaning, and we therefore profiled early weaning (postnatal day (P) 15), weaning (P20), post-weaning (P25), and adult (N=45-60 mice per age) mice to investigate foraging pattern development, as well as parental and genetic effects. Our design uses reciprocal crosses of *Mus musculus castaneous* (Cast) and C57BL6/J (B6) mice, which are wild-versus lab-derived subspecies of mice separated by ~500,000 years (Zheng et al., 2014). They are phenotypically distinct (see Table S2). By comparing the F1cb (Cast mother x B6 father) and F1bc (B6 mother X Cast father) hybrid offspring from these crosses, we test how parental effects influence foraging (Fig. 2A). The parental effects, as previously described (Lacey, 1998), potentially encompass *environmental parental effects* related to the in utero environment or parental behaviors, and *genetic parental effects* involving changes to imprinted genes, X-linked genes or mitochondrial genes that are preferentially inherited from one parent. Genetic effects due to differences between Cast and B6 alleles are uncovered by comparing F1bc offspring to B6 offspring, or F1cb offspring to Cast offspring (Fig. 2A, N= 15-20). This allows us to enrich for genetic effects because the mice share the same *maternal effects*, but are genetically different (Fig. 2A), though possible paternal influences are not eliminated.

At P15, only ~20% of B6 offspring entered the arena from the home base, while over 80% of F1cb, F1bc and Cast offspring entered during both phases of the assay (Fig. 2-S1A,B). However, only ~20% of F1cb, F1bc and Cast P15 animals consumed seeds in the assay. Thus, the full repertoire of foraging behavior in our assay has not developed at P15, but early stage patterns are captured from Cast, F1cb and F1bc offspring (Fig. 2-S1B). By P20, all mice of all crosses enter the arena, consume seeds and display independent foraging.

We profiled foraging for 191 mice across the different ages (P15, P20, P25, Adult), genetic backgrounds and crosses (B6, F1cb, F1bc and Cast), and captured measures for the feature set introduced above (Fig. 1-S1, Table S3). Distinguishing patterns shaped by parental or genetic effects at each age were revealed using Random Forest machine learning to trained classifiers that detect mice according to parental cross (F1cb vs F1bc) or genetic background (Cast vs F1cb; B6 vs F1bc). The results revealed F1cb and F1bc mice are not significantly distinguishable at P15 or P20, indicating parental effects at weaning are undetectable from our feature set (Fig. 2B, Parental Effects). This observation is intriguing because weaning is the stage during which the bond between mother and offspring changes and one might have predicted the opposite result (Lee, 1996; Trivers, 1974). Instead, we found that significant parental effects emerge *after* weaning. Both P25 and adult F1cb and F1bc offspring are differentiated greater than chance based on their foraging patterns (Fig. 2B). For example, F1cb and F1bc adults are distinguished with 67% sensitivity and 88% specificity (P=0.002, Fisher’s Test of Confusion Matrix). The OOB error is 23%, showing the classifier generalizes to a majority of cases (77%) (Fig. 2B).

While parental effects manifested at later ages, genetic effects robustly shaped foraging patterns at all ages beginning from P15, the earliest age tested (Fig. 2B). Cast patterns are distinguished from F1cb (and F1bc) patterns at P15 with 92% specificity and 100% sensitivity (Fig. 2B, Genetic Effects). B6 mice do not forage at P15 and are therefore different from F1bc (and F1cb) offspring due to genetic effects (Fig. 1-S2). Differences persisted across older ages and B6 mice were distinguished from F1bc (and F1cb) offspring with 100% specificity and sensitivity at P20, P25 and in adults (Fig. 2B). These data show that parental and genetic effects in Mus musculus have distinct development stages of influence on offspring foraging patterns. The next challenges include determining the nature of these patterns and whether they involve modules.

### Parental and Genetic Effects Shape Foraging Patterns Differently at Different Ages

The nature of the foraging patterns exhibited by the mice above can be extracted from the random forest classifiers we trained to detect significant parental or genetic effects. In one outcome, parental and/or genetic effects shape similar patterns across the different significantly impacted ages. Alternatively, parental and/or genetic effects manifest differently at different ages. To learn which occurs, we defined the informative features for the distinguishing patterns identified by the random forest classifiers for parental or genetic effects at each age. Informative features were identified, and non-informative features eliminated, using permuted importance analysis, which normalizes random forest feature importance measures, corrects feature importance bias and identifies significantly informative features in a classifier (Altmann et al., 2010). We found that significant features characterizing parental effects on P25 patterns are completely different from those characterizing effects on adult patterns (Fig. 2C, Permutated Importance values, P<0.01). Thus, parental effects shape foraging patterns differently at different ages.

The patterns of parental effects were visualized using radar plots of the data for significantly informative features (Fig. 2D and E) (see Table S1 for feature definitions, Table S3 for data). At P25, a top ranked feature is total visits to Pot#2 during the Exploration phase (E.TVP2.0), which is greater in F1cb than F1bc offspring (Fig. 2D). In contrast, a top feature distinguishing adult F1cb and F1bc foraging is the total visits to the center zone during the third time bin of the foraging phase (F.TVC.3) (Fig. 2E). F1cb mice perform this behavior more frequently, while F1bc mice spend more total time at the food patch at Pot#4 during the same time period (F.TTP4.3) (Fig. 2E). The results show parental effects shape complex, non-obvious patterns that differ according to age.

Intriguingly, genetic effects also shape foraging differently at different ages (Fig. 2-S2). A Venn diagram of the features with significant importance values in the P20, P25 and adult classifiers for genetic effects shows that none of the features are common to all ages (Fig. 2-S2A). Further, the majority of significant features are age-specific (Fig. 2-S2A). Radar plots show the distinguishing patterns due to genetic effects at each age (Fig. 2-S2B-D). For example, a feature distinguishing B6 from F1bc (and F1cb) mice at P20 is the total time in sector 2 during the last time bin of the Exploration phase (E.TTS2.5), which is highest in B6 (Fig. 2-S2B). A significant feature in adults, but not other ages, is the peak velocity during the Exploration phase (E.PeakVelocity), which is higher in F1bc/F1cb mice compared to B6 (Fig. 2-S2D). The basis of these age-dependent patterns could involve changes to specific modules, which we investigate next.

### Discovery of 59 Discrete Modules Underlying Foraging Across Different Ages, Crosses and Genetic Backgrounds

To test whether modules underlie foraging patterns, DeepFeat was applied to the XY tracking data for the Cast, F1cb, F1bc and B6 mice in our cohort. We found 6,585 home base excursions, and unsupervised hierarchical clustering and dynamic tree cutting identified 59 distinct behavioral modules from these excursions (Fig. 3A). If the modules detect meaningful behavioral sequences, they are expected to be sensitive to context and age. We quantified the expression frequency of each module for each mouse in each phase of the assay and display the results as a heatmap showing mean expression frequencies according to phase and age (Fig. 3B). Both phase (context) and age dependent differences are apparent from the module expression data and clustering grouped sets of modules with similar patterns (Fig. 3B). Chi-Square tests of independence confirmed that module expression frequencies depend significantly on the phase (P= 2.3×10^−85^), and on age (P=7.0×10^−64^). We conclude that the modules are bona fide, biological behavioral sequences. Support for this claim grows further below.

**Figure 3.**
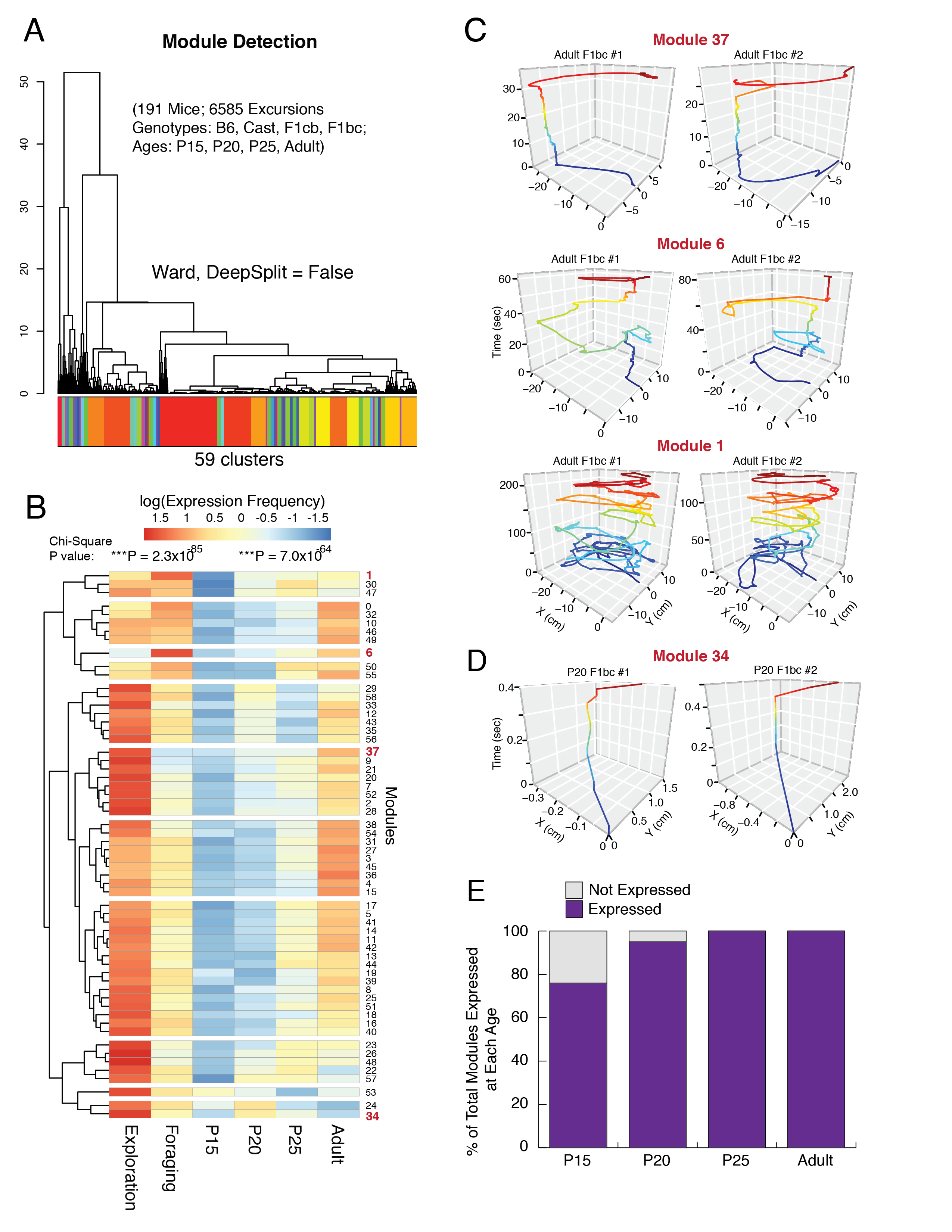
Identification of 59 behavioral modules underlying mouse foraging patterns across different ages reveals contextual and developmental changes to module expression. **(A)** Dendrogram of the clustering results to define modules from home base excursions. DeepFeat identified 59 modules from over 6500 home base excursions performed by 191 mice in our cohort of B6, Cast, F1cb and F1bc offspring from four ages. Distinct clusters (modules) are indicated by color. **(B)** The heatmap shows the relative expression frequency of different modules by phase (Exploration versus Foraging) and age. The module expression counts are centered by row. Clustering on the rows identified 11 groups based on expression frequency similarities. P values for a Chi-Square test of independence are shown for Phase and Age on the top of the chart. The results show that module expression patterns are significantly dependent on phase and age. **(C and D)** Examples of representative modules from two different adult F1bc mice for Modules 37, 6 and 1 (C), and for two different P20 F1bc mice for Module 34 (D). The data show the XY tracking pattern of the mice over time during one round trip excursion from the tunnel to the home base cage (0,0). Behavioral sequences are colored according to time (normalized to total time). **(E)** Bar plot shows the percentage of the 59 modules expressed (purple) by mice according to age. Modules not expressed (grey) have a frequency ≤ 1 across all mice tested for that age.

Our data indicate increased module expression frequency and diversity in the Exploration compared to Foraging phase, revealing contextual effects on module expression (Fig. 3B). One example of a module that is especially enriched in the Exploration phase is Module 37, which involves a direct round trip from the home base to the food patch (Pot#2) (Fig. 3B, C). We found two modules are expressed more frequently in the Foraging phase, including Modules 6 and 1 (Fig. 3B,C). Module 6 involves a complex pattern with a visit to the new food patch in the Foraging phase (Pot#4) followed by a looping survey of the former patch (Pot#2) (Fig. 3C). In contrast, Module 1 involves a long, extended survey of the environment without an obvious destination (Fig. 3C). Nine modules are expressed similarly in the Exploration and Foraging phases, suggesting context independence in terms of expression frequency (Modules: 30, 47, 0, 32, 10, 46, 49, 50 and 55) (Fig. 3B).

Developmental changes to the expression of specific modules are revealed in the data. For example, Module 37 is predominantly expressed by adults (Fig. 3B,C), while Module 34, which involves a sub-second dart into and out of the arena, is predominantly expressed by P20 juveniles (Fig. 3B,D). All 59 modules are expressed by adults and P25 juveniles at some frequency, but 14% and 5% of modules are not expressed by P15 and P20 weanlings, respectively (expression frequency ≤ 1 across all mice) (Fig. 3E). Modules 50 and 55, for example, emerge after P20. Thus, our data reveal modules underlying foraging, show changes to module expression in naïve versus more goal orientated contexts and uncover a time course for the development of discrete modules.

### Genetic Effects in Mice Shape Different Foraging Patterns Through Changes to the Expression of Shared Modules, Not the Creation of Novel Modules

The genetic effects that shape different foraging patterns in Cast, B6 and F1bc/F1cb mice (Fig. 2) could involve the creation of distinct modules in one strain versus another and/or changes to the expression of a shared set of core modules. Here, we set out to determine which occurs and uncover a basis for genetic differences in Mus musculus foraging patterns. Our analysis of the DeepFeat output did not find any modules that are solely expressed by one genetic strain, but not another (eg. B6, but not F1bc). This indicates the genetic differences do not give rise to different modules, and the 59 modules we uncovered are shared across all strains. Instead, genetic differences in foraging might involve changes to the expression of specific modules.

To test our prediction, random forest machine learning was used to train classifiers to detect Cast, B6 and F1 hybrid (F1bc/F1bc) mice at each age based on module expression data (Fig. 4A). We found that each genotype was detected at rates significantly better than chance at each age, with an average of 35% sensitivity and 99-100% specificity (Fig. 4A). The OOB error was low in most cases (≤10%) (Fig. 4A), indicating highly generalizable models. From the classifiers, we uncovered significantly informative modules from permuted importance values (P<0.05). The results reveal discrete modules with distinguishing expression patterns for each genetic strain at each age.

**Figure 4.**
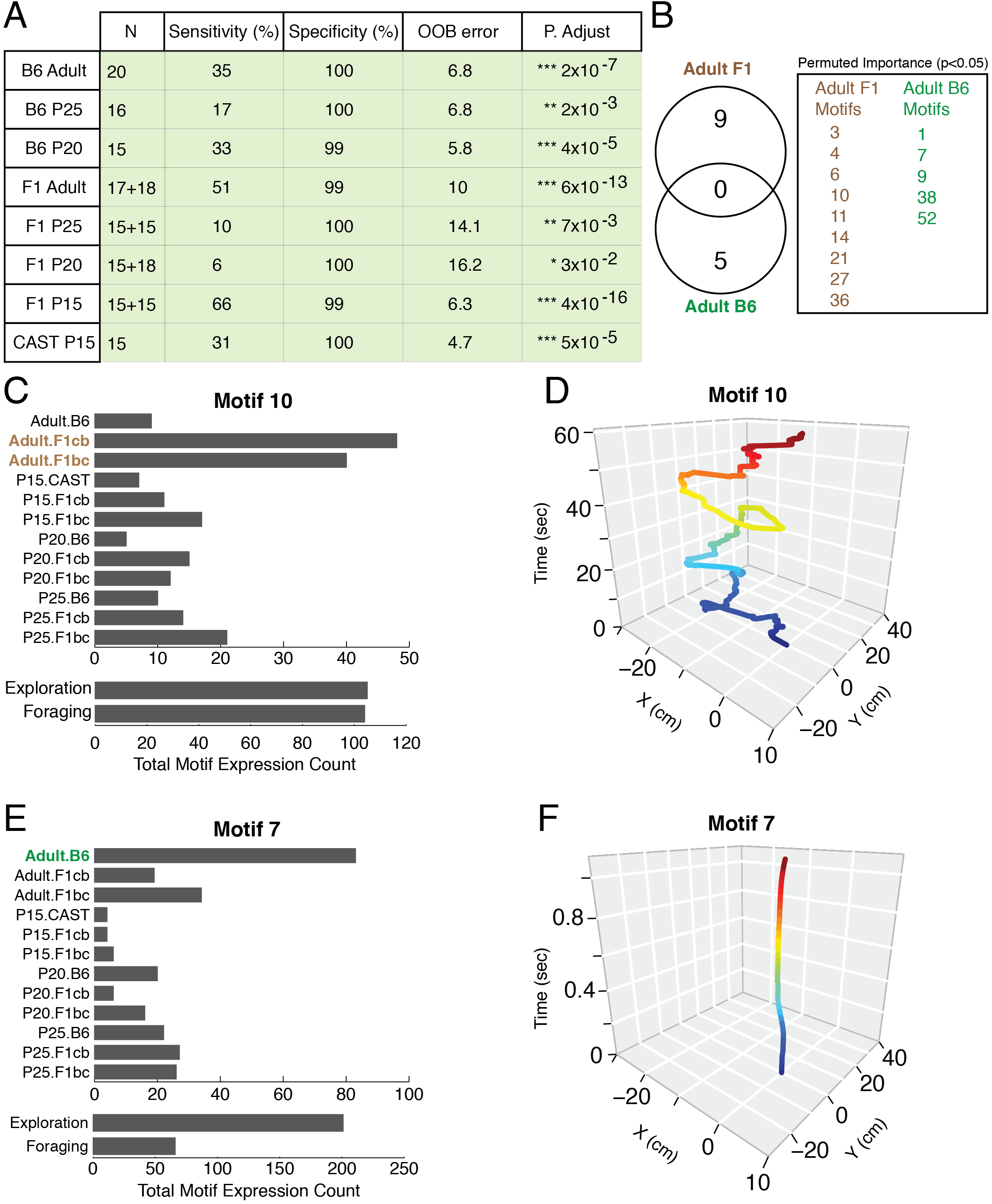
Changes to the expression of specific behavioral modules underlie genetic effects on foraging patterns. **(A)** The table shows the results of the random forest classification of animals by genotype (B6, F1 hybrid and Cast) for each age (P15, P20, P25 and adult). A two-tailed Fisher’s test on the resulting confusion matrix showed statistically significant detection of each group (Benjamini-Hochberg adjusted p-values shown). OOB, out-of-bag error. N shows number of animals tested (F1 hybrid: F1cb + F1bc). **(B)** Permuted importance analysis for the classifiers trained to detect adult F1 hybrid (F1cb and F1bc) and B6 mice identified significantly informative modules (P<0.05). The Venn diagram and box inset show the modules informative for adult B6 versus F1 hybrid mice phenotypes, and they are different (refer to Figure 2B for context and age expression details). **(C and D)** Module 10 is a significantly informative module discriminating F1 hybrid adult behavior. Counts of the module’s expression frequency show it is expressed by all strains and ages, but is relatively highly expressed by F1cb and F1bc adults (C), and equally in the Exploration and Foraging phases. A representative trace of the module is shown from an F1bc adult, indicating a long, looping trajectory with multiple pauses in different locations on the side of the arena harboring Pots 1 and 2 (D). **(E and F)** Module 7 is a significantly informative module discriminating B6 adults. It is expressed highly by B6 adults, and more frequently in the Exploration compared to the Foraging phase (E). A representative adult B6 trace reveals that Module 7 is a brief, sub-second emergence from the tunnel to the threshold of the arena and back (F).

For example, adult F1 hybrids (F1bc and F1cb) are distinguished from all other strains and ages by the expression of 9 modules (Fig. 4B). Module 10 is one example and is most frequently expressed by F1cb and F1bc adults, though others also express it (Fig. 4C,D). This module is relatively insensitive to context and equally expressed in the Exploration and Foraging phases (Fig. 4C,D). B6 adults are distinguished from all other strains and ages by differences in the expression of 5 modules. Module 7 is one example and involves a brief entry and pause on the platform (Fig. 4E,F). It is most frequently expressed by B6 adults and shows context dependent expression with enrichment in the Exploration phase (Fig. 4E,F). Thus, genetic differences in mice cause changes to the expression of specific modules underlying foraging patterns.

Finally, in our analysis we found that some modules appear visually similar, raising the concern that they are not truly distinct module subtypes. We therefore performed a more detailed analysis. For instance, Modules 7 (Fig. 4F), 34 (Fig. 3D), 54 and 30 (Fig. 4-S1A), are short, similar looking sequences expressed near the tunnel to the home base cage. However, when we compared descriptive statistics for the phenovectors of excursions classified under these four module types, differences became apparent. Module 34, for example, is expressed more frequently during the middle of the Foraging phase, while the others are expressed more frequently at the beginning (Fig. 4-S1B). Modules 7, 30, 34 and 54 have differences in the kurtosis of the XY movement patterns and/or different lengths (number of frames) (Fig. 4-S1C-E). Modules 7 and 54 differ in terms of the average range of movement in the Y axis (Fig. 4-S1H,I). Further, all four modules have different expression patterns by phase, age and genotype (Fig. 4-S1J-L). Module 7 is uniquely highly expressed during the Exploration phase and by adults of the B6 genotype (Fig. 4-S1J-L). Module 34 is expressed predominantly by F1cb and F1bc P20 juveniles in the Exploration phase (Fig. 4-S1J-L). We conclude that these modules are in fact different, and that brief, ~1 second behavioral sequences carry phenotypic information reflecting genotype, age and/or context. Overall, these insights, and the observed genetic effects from distant subspecies of mice, help reveal how evolution shapes foraging through modules.

### Different Modules Link to Memory, Reward, Risk and Effort Responses

Different modules are expected to have different functions. If this is true, specific modules should relate to specific keystone features of foraging that have clear ecological functions. Food consumption is one keystone feature that is an ethologically relevant measure of reward and motivation. Thus, finding modules significantly associated with the total amount of seeds eaten in our assay (TFC.0, see Table S1) could reveal putative modules for reward, feeding and food patch exploitation. A keystone feature for risk-related behavior is thigmotaxis, which is an evolutionarily conserved strategy for predator and risk avoidance that involves aversion to exposure and increased proximity to walls (Prut and Belzung, 2003; Walz et al., 2016). Modules significantly associated with the percentage of time spent in the exposed center zone of the arena (PTC.0) are therefore putative “risk” behavior modules (Fig. 1-S1A and Table S1). Two keystone features for effort and energy expenditure are the total amount of sand dug from the pots (TSD.0) and the total distance traveled (TD, see Table S1), potentially allowing the identification of putative foraging “effort” modules. Finally, our assay includes reversal learning and perseveration, providing an enticing opportunity to uncover modules for memory and food patch abandonment responses. The reversal learning and perseveration involve the following.

During the Exploration phase, mice learn that Pot#2 harbors food (Movie S1) and spend significantly more time at this Pot (Fig. 5-S1A). During the Foraging phase, they learn to abandon this now absent food source and discover the hidden food in Pot#4 (Movie S2). However, mice spend time searching the former food source, especially during early stages of the Foraging phase (Fig. 5A and 5-S1A) and excavate much of the Pot#2 sand during their search (Movie S2). The amount of Pot#2 sand dug (F.TSDP2) is therefore one keystone feature of the perseverative effects of the learned memory.

**Figure 5.**
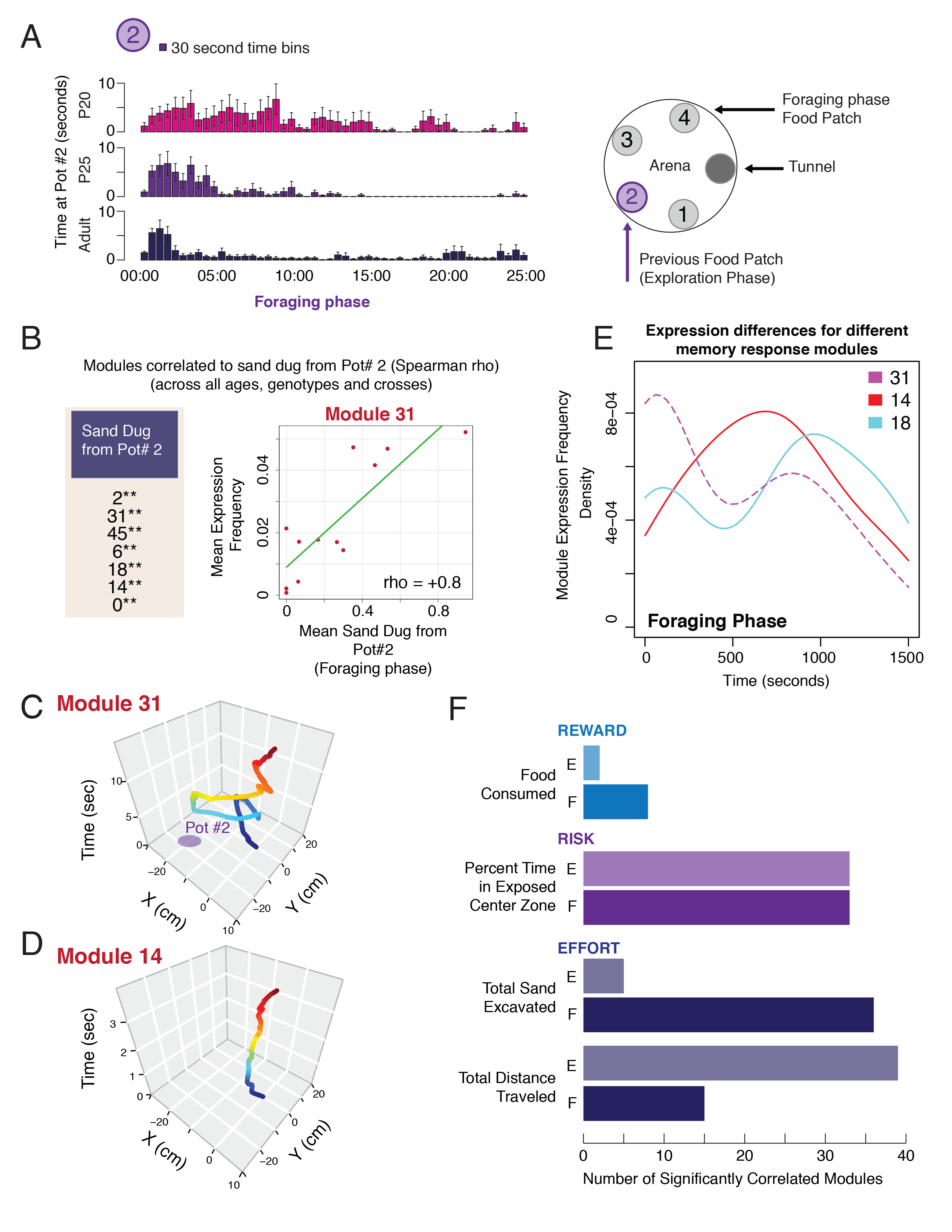
Different modules are linked memory, reward, risk and effort responses during foraging. **(A)** P20, P25 and adult mice spend increased time investigating the former food pot (Pot#2) during the Foraging phase, showing perseverative effects of the learned food location. The bar chart shows time at Pot#2 over the course of the 25 minute Foraging phase trial for B6 mice in 30 second time bins. The error bars are SEM. The schematic indicates the layout of the arena and location of Pot#2. **(B)** Top putative memory and perseveration response modules are revealed from significant correlations to the amount of sand dug from Pot#2 in the Foraging phase are shown (P<0.01, Spearman Rank correlation). An example of the data is shown for Module 31. The average expression frequency for the module across the 12 different genotype, cross and age groups is correlated to the average sand dug from Pot#2 in the Foraging phase. **(C and D)** Representative traces for the memory and perseveration Modules 31 and 14. Both modules are significantly associated with the amount of sand dug from Pot#2, but have distinctive temporal and movement patterns. Module 31 is longer and involves a looping visit to Pot#2, while Module 14 is a brief, multi-second emergence from the tunnel and back. **(E)** The density plots show the expression start times for all Foraging phase excursions classified as the memory and perseveration Modules 31, 6, 14 or 18. The data reveals differences in the temporal expression patterns of the modules across the entire 25 minute Foraging phase trial (see text). **(F)** The bar chart shows the number of modules significantly correlated to reward (blue, total food consumed), risk (purple, percent time in center zone) and effort (dark blue, total sand dug and total distance traveled) keystone features in the Exploration (E) and Foraging (F) phases. Spearman rank correlation P<0.05.

To identify putative functionally distinct modules linked to reward (TFC.0), risk (PTC.0), effort (TSD and TD) and memory (F.TSDP2) responses, we calculated the mean values for each keystone feature across mice in the 12 different ages, genotypes and crosses in our cohort, and tested whether specific modules are significantly correlated to specific keystone features (Spearman rank correlation). For memory and perseverative responses, we identified 15 modules that are significantly correlated to the amount of sand dug from Pot#2 during the Foraging phase (Spearman rho, P<0.05). For example, a plot contrasting the average Foraging phase sand dug from Pot#2 to the average frequency of Module 31 reveals a significant positive correlation (Fig. 5B, rho = +0.8, P = 0.004). Examination of the Module 31 structure during the Foraging phase shows a visit to Pot#2 in the middle of excursion (Fig. 5C). However, other putative memory response modules have different structures, as well as temporal patterns of expression. For example, Module 14 is a relatively short excursion sequence performed at the tunnel to the home base (Fig. 5D). While Module 31 is frequently expressed during early and late stages of the Foraging phase (Fig. 5E), Module 14 is predominantly expressed during middle stages (Fig. 5E). Another case, Module 18 (Fig. 5E), is highly expressed during the late stages of the phase (Fig. 5C). We conclude that discrete modules are linked to memory and perseverative responses during foraging and show they are expressed at different times, possibly reflecting changes to internal state and adaptive processes for food patch abandonment.

Using a similar approach, we uncovered 8 modules significantly linked to food consumption (reward) in the Exploration and/or Foraging phase contexts of the assay (Fig. 5F). For example, Module 27 is highly correlated to food consumption (rho = +0.7, P=0.009) and involves extended stops at Pot#4, the Foraging phase food patch (Fig. 5-S1B). We found 35 modules linked to risk (Fig. 5F), including Module 2, which is a high risk module involving many traverses of the exposed center and a relatively long 1-2 minute excursion from home (Fig. 5-S1C; rho = +0.9, P= 3×10^−5^). Finally, 36 and 40 putative “effort” modules are significantly linked to the total sand dug from all Pots and the total distance traveled, respectively (Fig. 5F). One example of a top module linked to distance traveled is Module 19 (rho = +0.9, P=2×10^−5^), which involves a long, looping ~1 minute excursion from the home (Fig. 5-S1D). Therefore, distinct modules are linked to distinct ecologically important elements of foraging, suggestive of functional differences.

### Specific Behavioral Modules, Genes and Pathways are Sensitive to Parental Effects in Offspring

Our studies of foraging patterns revealed that parental effects on foraging increase with age (Fig. 2). In particular, we observed the strongest phenotypic effects in adults, but no effects in P15 and P20 weanlings. Here, we seek to test whether parental effects impact the expression of specific modules in adult offspring and identify candidate genetic mechanisms for shaping foraging during later stages of life. To define modules impacted by parental effects in adult offspring, we trained a random forest classifier to detect F1cb from F1bc adults from module expression data. Significant detection of F1cb from F1bc adults (P=0.0005) was observed with 88% sensitivity, 72% specificity and a 20% OOB generalization error (Fig. 6A). Permuted importance values for modules in the classifier revealed five significantly informative modules (P<0.05) (Modules: 1, 6, 21, 24 and 40). Bar plots of the expression frequencies for the significant modules show how the F1cb versus F1bc adults differentially express these modules during foraging (Fig. 6B). Having identified discrete modules impacted by parental effects in adults, we set out to uncover mechanisms involved.

**Figure 6.**
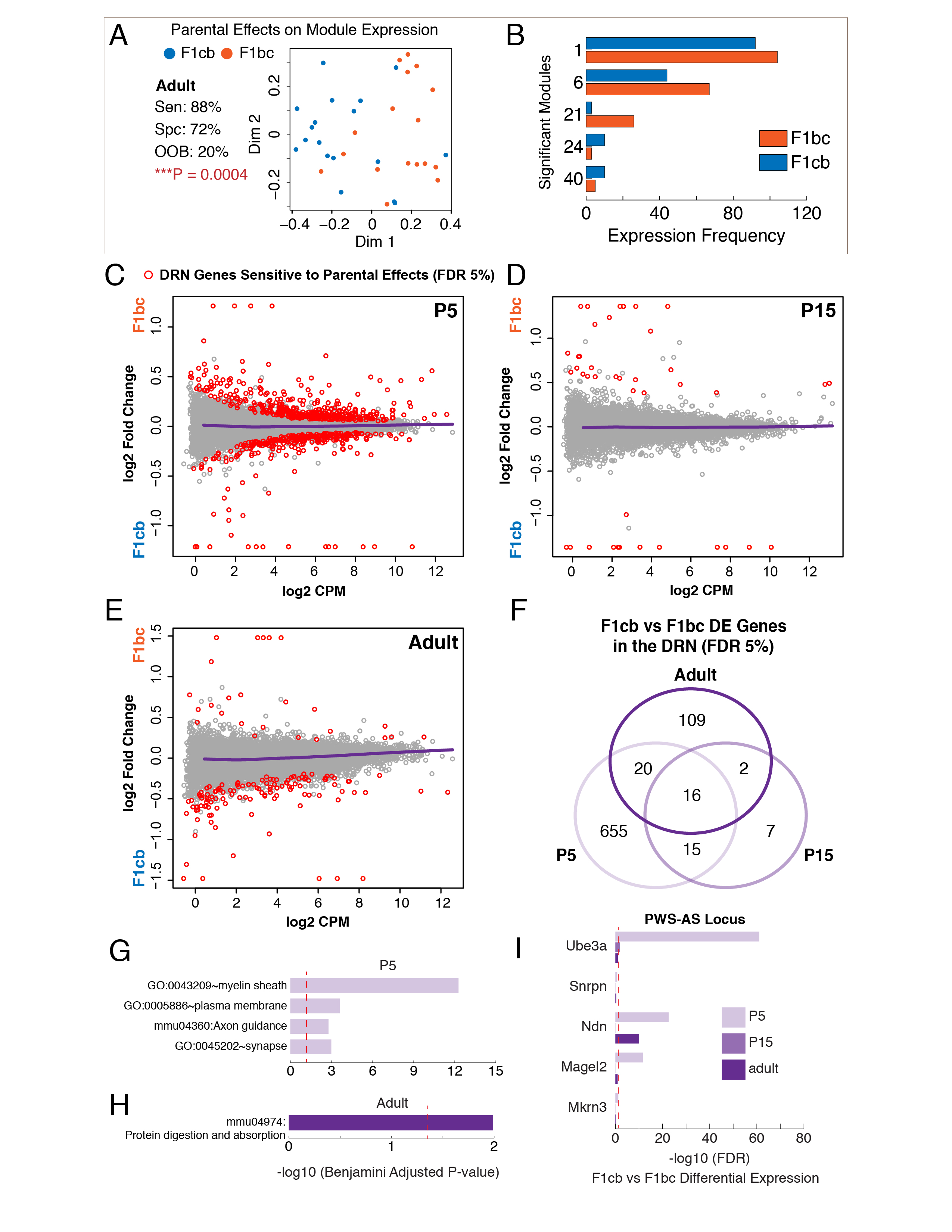
Discrete behavioral modules, genes and pathways are impacted by parental effects in offspring in an age-dependent manner. **(A)** Multi-dimensional scaling plot of random forest classification results for adult F1cb versus F1bc mice based on module expression data. The results show significant discrimination by the trained classifier. **(B)** The bar plots show the adult F1cb and F1bc expression frequency data for significantly informative modules in the trained classifier. Permuted Importance Analysis (P<0.05). **(C-E)** MA-plots of RNASeq analysis of differential gene expression in the DRN region of F1cb versus F1bc offspring at P5 (C, N=7), P15 (D, N=7) and in adults (E, N=8). Significant genes are shown in red (5% FDR). The data show a relative reduction of parental effects in P15 juveniles, consistent with our analysis of behavior patterns (Fig. 2B). **(F)** Venn diagram of P5, P15 and adult differentially expressed (DE) genes due to parental effects. **(G and H)** Bar plots show the results of DAVID gene ontology and pathway enrichment analysis for genes with significant parental effects in the P5 (G) and adult (H) DRN. Only statistically significant enrichment terms are shown (5% FDR threshold indicated by red line). The x-axis shows the adjusted p-value for the ontology term enrichment. **(I)** Bar chart of the adjusted p-values for DRN differential expression between F1cb and F1bc offspring for imprinted genes in PWS-AS locus. *Ube3a, Magel2* and *Ndn* show significant expression level differences at P5. *Ndn* is also significant in adults and *Ube3a* is significant at P15. Red line indicates the threshold for a 5% FDR (see Table S?).

To identify candidate genes and pathways in the brain impacted by parental effects, we tested for gene expression differences between F1cb and F1bc female offspring during nursing (P5), weaning (P15) and in adulthood (Fig. 6C-E). Our study focuses on the dorsal raphe nucleus (DRN) region of the midbrain, which contains the largest serotonergic nucleus innervating the forebrain. This region exhibits developmental changes that impact feeding and activity, is linked to mental disorders, and modulates diverse systems involved in foraging, including reward, feeding, stress & anxiety, exploratory drive, arousal and sensorimotor systems (Bromberg-Martin et al., 2010; Jean et al., 2017; Lowry et al., 2008; Monti, 2010; Nectow et al., 2017; Ranade and Mainen, 2009; Rood et al., 2014). We use RNASeq datasets we previously published (Huang et al., 2017) and compare DRN region gene expression in female F1cb to F1bc offspring for P5 nursing pups (N=7 for each cross), P15 weaning juveniles (N=7 for each cross) and 8-week old adults (N=9 for each cross). Genes subject to parental effects are significantly differentially expressed between F1cb versus F1bc offspring at a 5% false discovery rate (FDR) threshold using edgeR (Robinson et al., 2010) (Fig. 6C-F, Table S4).

At P5, 706 genes are significantly differentially expressed between F1cb and F1bc offspring, indicating parental effects (Fig. 6C,F). By P15, the number drops to only 40 genes, which is a 18-fold decrease compared to P5 (Fig. 6D,F). In adults, however, the number rebounds to 147 differentially expressed genes - a 4-fold increase relative to P15 (Fig. 6E,F). Importantly, the reduction in parental effects on gene expression at P15, but subsequent increase in adulthood, is concordant with the pattern of parental effects we observed at the level of foraging behavior (see Fig. 2). The results further reveal that the strongest parental effects on DRN gene expression manifest at P5, during nursing.

Genes differentially expressed between P5 F1cb and F1bc mice are enriched for *myelin sheath*, *plasma membrane, axon guidance* and *synapse* gene categories (Fig. 6G, Table S5). Changes to myelination, axon guidance and synapse function could shape offspring brain function and behavior patterns. No enrichments were observed from P15 differentially expressed genes. For the adult, we found enrichments for the KEGG *protein digestion and absorption* pathway, which is related to enrichments for ion channels (eg. *Kcnq1*, a voltage gated potassium channel) and several collagen genes (eg. *Col4a3*, collagen type IV alpha 3) (Fig. 6H). Our results reveal candidate genes and pathways involved in the parental effects shaping offspring foraging.

Among the most statistically significant candidate genes, we found several genes in the PWS – Angelman Syndrome (AS) locus, including *Ube3a, Magel2* and *Ndn* (Fig. 6I, Table S4). PWS patients have profoundly altered feeding, activity and motivated behaviors, and show developmental changes to their phenotypes (Cassidy et al., 2012). Mouse models also show altered motivation, anxiety, activity and feeding (Bervini and Herzog, 2013). Thus, these genes are strong candidates for shaping modules of economic behavior for foraging. By focusing on one of these genes, *Magel2*, we tested for age-dependent effects on foraging patterns and module expression.

### Loss of the PWS-AS Gene, *Magel2*, Shapes Adult Foraging Patterns by Modifying the Expression of Specific Behavioral Modules

*Magel2* influences circadian rhythm, feeding, anxiety, arousal and serotonin signaling in mice (Kozlov et al., 2007; Mercer et al., 2009). Roles in shaping more complex, ethological foraging patterns have not been examined. We bred published *Magel2* mutant mice to generate *Magel2^+/−^, Magel2^−/+^* and *Magel2^+/+^* female offspring. Since *Magel2* is imprinted and expressed from the paternal allele, *Magel2^+/−^* offspring are mutants compared to the other genotypes. Based on our results above, *Magel2* is a candidate mechanism for shaping foraging at later ages, after weaning. To test this, we compared foraging patterns between *Magel2^+/−^* mutants and controls at weaning (P20) and again in adults (9-12 weeks of age) (Table S6).

Random forest machine learning on the data captured for foraging features did not uncover a significantly distinguishing pattern in the *Magel2^+/−^* mutants at P20 (Fig. 7A, P=0.55). However, in adults, *Magel2^+/−^* mutants were identified with 86% accuracy, which is significantly better than chance (Fig. 7A, P=9.4×10^−7^). Therefore, loss of *Magel2* more strongly impacts adult compared to P20 foraging patterns, supporting our hypothesis. The informative features in the classifier detecting *Magel2^+/−^* mutants from controls included 10 Foraging phase features and 2 Exploration phase features (Fig. 7B). The top ranked feature is the total time in the arena during the Foraging phase (F.TTA.0, Table S1 and S6) and radar plots of the data show that *Magel2^+/−^* mutants spend less time in the arena during the foraging phase and more time in the tunnel entry (F.TTE.2) and tunnel zone (F.TTTZ.0), and perform more total visits to the tunnel during the first time bin (F.TVT.1) (Fig. 7B). Our next goal was to determine whether this distinguishing pattern of behavior involved changes to specific behavioral modules, either by giving rise to new mutant modules or changing the expression of core modules shared across mutants and controls.

**Figure 7.**
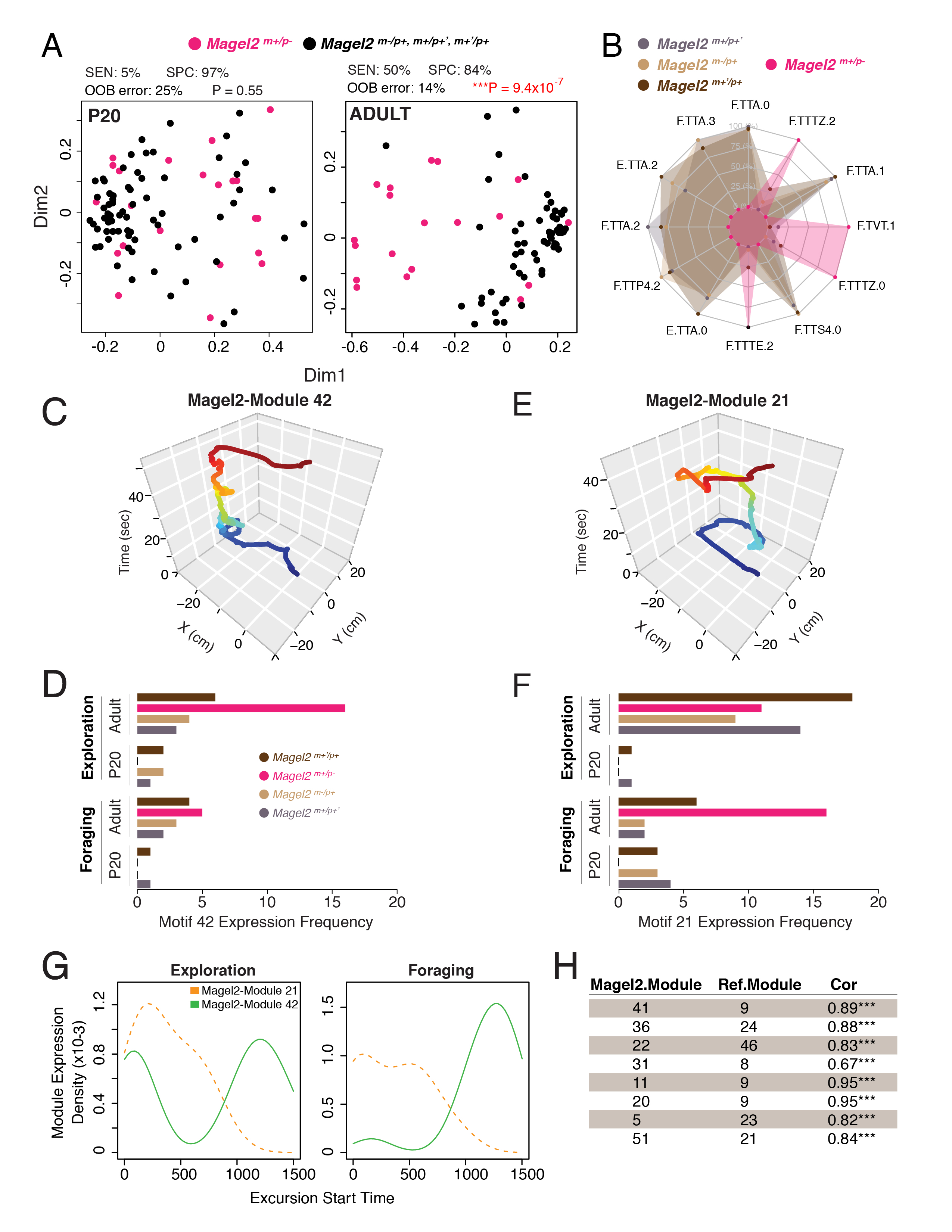
The PWS gene, *Magel2*, most strongly shapes adult behavioral patterns associated with the home. (**A**) Multidimensional scaling plots of the random forest classifier proximity values to detect *Magel2* mutants (*Magel2^m+/p−^*) from controls (*Magel2^m+/p+’^*, *Magel2^m-/p+^* and *Magel2^m+’/p+^*) at P20 and in adulthood. The sensitivity (SEN), specificity (SPC), out-of-bag error (OOB) and the two-tailed fisher test P-value for the confusion matrix of the classification results are shown above each plot. **(B)** Radar plot shows the relative values of data for the significantly informative features for the classifier detecting *Magel2^m+/p−^* adults (yellow) from *Magel2^m+/p+’^* (brown), *Magel2^m-/p+^* (green) and *Magel2^m+’/p+^* (pink) controls (P<0.01, PIMP). The results show the distinguishing pattern in *Magel2^m+/p−^* adults (see main text and Table S1 for feature descriptions). **(C-F)** A representative trace of Magel2-Module 42 (C) and Magel2-Module 21 (D), significantly informative modules in the adult *Magel2^m+/p−^* classifier. The bar plots show the expression frequency of the modules for all genotypes and ages (D and E). The results show phase (context) dependent changes to module expression in the mutants. **(G)** Density plots show distinct temporal expression patterns (excursion start times) across the 25 minute Exploration and Foraging phase trials for excursions classified as Module 42 versus Module 21, two module types significantly linked to phenotypic effects in *Magel2^m+/p−^* adult mutants. **(H)** The table shows the top reference module matches for each of the nine Magel2-Modules that are significantly informative for the *Magel2^m+/p−^* adult mutant phenotype. Magel2-Module and reference modules IDs are shown. Cor, is the Pearson correlation for the match of the eigenphenotype. ***Holmes adjusted p-value < 10^−10^.

To identify modules, DeepFeat was applied to analyze the 5,302 home base excursions detected from the P20 and adult *Magel2* mutant and control mice. This initial analysis identified 42 modules. However, random forest classification based on the expression *frequency* of these modules was unable to distinguish adult mutants from controls better than chance (P>0.05, Fisher’s test of the confusion matrix). We therefore concluded that loss of *Magel2* does not create new modules in mutants compared to the controls, and changes to the expression frequency of modules is not a significant component of the mutant phenotype. Alternatively, changes to the *contextual pattern* of module expression might be involved.

In a revised analysis, we integrated excursion start time and phase information into the phenovectors captured by DeepFeat. With these new elements added to the XY movements, DeepFeat now identified 75 different modules from the 5,302 home base excursions (Fig. 7-S1A, Table S6). These modules yielded statistically significant detection of the adult mutant mice (Fig. 7-S1B). Nine modules were identified as significantly informative in the adult mutant classifier (Importance permutation P<0.05) (Fig. 7-S1C). Magel2-Module_42 is the top ranked module and involves an extended visit near Pot#2 (Fig. 7C). This module is expressed relatively equally in Exploration and Foraging phases by the controls. However, adult mutants exhibit increased expression during the Exploration phase (Fig. 7D), consistent with a context-dependent shift.

Magel2-Module_21 is another significantly informative module for the mutant phenotype, and involves prolonged lingering near the tunnel to the home base (Fig. 7E). In controls, this module is preferentially expressed in the Exploration phase (Fig. 7F). However, adult mutants exhibit increased expression during the Foraging phase (Fig. 7F). The altered contextual expression of this module is consistent with many aspects of the mutant’s foraging pattern, including increased time in the tunnel zone during the Foraging phase (F.TTTZ) (Fig. 7B). Temporally, Magel2-Modules 42 and 21 have distinct expression patterns, indicating roles at different stages of foraging behavior (Fig. 7G). Overall, the basis of the phenotype in adult *Magel2* mutants is revealed to involve context- and age-dependent changes to the expression of nine discrete modules.

Finally, we sought to test whether the modules impacted by loss of *Magel2* related to the modules impacted by parental effects in adult mice (Fig. 6B). To compare are modules across different cohorts of mice, we developed a module matching strategy based on ideas for comparing gene networks (Langfelder and Horvath, 2007; Langfelder et al., 2011). From the phenovector data for each of the 59 reference modules identified in Cast, B6, F1cb and F1bc mice, we computed the first principal component for each module. This eigenvector reflects the major phenotypic characteristics of a module, which we call the eigenphenotype. We then computed the eigenphenotypes for the nine modules informative for the adult *Magel2* mutant phenotype after removing the start time and phase information, so that the underlying data matches the reference modules. The Magel2-Modules were matched to the reference modules using a Pearson correlation of the eigenphenotypes (Fig. 7-S2). The results show the top significant module matches, revealing how the *Magel2* modules map to specific reference modules (Fig. 7H). Importantly, we found that Magel2-Modules 36 and 51 match to reference modules 24 and 21 (Fig. 7H), respectively, which are two of the five modules impacted by parental effects in adult F1cb versus F1bc mice (Fig. 6B). Reference module 21 is a significant “risk” module (E.PTC: P=0.005, F.PTC: P=0.01) and also linked to effort in the Foraging phase (F.TSD: P=0.002) (see Fig. 5F). The results validate *Magel2* as a mechanism shaping the expression of specific modules during later life, and establish a framework for phenotypic comparisons across studies.

## DISCUSSION

We set out to learn whether foraging patterns are built from discrete modules, and behavior paradigm with custom analysis methods, we found an atlas of 59 core modules of economic behavior. Each module reflects a type of behavioral sequence performed during round trips from the home. Distinct modules are linked to reward, risk, effort and memory responses during foraging, indicating putative functional subclasses and relationships to keystone features of foraging patterns. Mice were found to express different modules in different temporal sequences that depended on context. The modular architecture of foraging increases in complexity over development and we defined ages when specific modules emerge. Intriguingly, parental and genetic effects in mice are revealed to shape foraging in a highly age-dependent manner by changing the expression of core modules, rather than creating new modules. Parental effects impact foraging increasingly as mice age and we identify candidate genes and pathways involved. Validation of one gene, *Magel2*, confirmed increased phenotypic effects in adults compared to juveniles. Further, loss of *Magel2* function, which causes PWS and autism in humans, alters foraging by changing the contextual expression of nine modules in adults. A subset of these modules matched to the modules sensitive to parental effects, revealing a putative mechanistic link through *Magel2*. Our results help elucidate the basis of normal and abnormal behaviors, and how evolution shapes new adaptive patterns.

### Towards Understanding How the Genome Shapes Complex Economic Behavior Patterns

Most neuroscience studies use controlled tests aimed at specific behavioral responses, rather than studying rich behavior patterns. As a consequence, the principles and importance of deeply analyzing relatively spontaneous behavioral sequences in rodents was emphasized decades ago (Kavanau, 1967). More recently, sub-second patterns of body language were uncovered using machine learning in mice (Wiltschko et al., 2015), revealing an important new dimension of behavior. Here, we also identified important phenotypic information from short behavioral modules (1-2 seconds or less), supporting the rationale for further studies in this area. Invertebrate model organisms have shown how computational dissections of behavioral sequences can ultimately help resolve links between specific mechanisms and behavior patterns (Robie et al., 2017; Vogelstein et al., 2014), and advances are being made in mammals (Hong et al., 2015; Kabra et al., 2012; Markowitz et al., 2018; Weissbrod et al., 2013). However, the study of modularity in free patterns of complex ethological behaviors presents many challenges in terms of how to design the assay, select features for analysis and analyze the data to uncover biologically important patterns and modules. Our study makes advances on these fronts for foraging and related economic behaviors.

We found that modularity in foraging emerges from behavioral sequences anchored by the home base. The role of the home base in behavior is understudied, with the exception of a few pioneering efforts (Eilam and Golani, 1989; Gorny et al., 2002). However, the centrality of the home in the ecology of many species, including humans, is undeniable. Humans spend 69% of their life in their home (Klepeis et al., 2001) and homelessness is a major factor associated with poverty, mental illness and addiction, yet we know little about the role of the home in structuring adult and developing behavior patterns. For many species in the wild, behavioral responses are structured within a defined home range (Börger et al., 2008). Neural mechanisms for navigating and dead reckoning relative to a landmark, such as the home, have been identified (Wallace et al., 2008). Thus, the modular structure of foraging that we uncovered is likely to have been selected for, having a genetic basis and analogs across different species. In support of this, we show that genetic differences between distant strains of *Mus musculus* shape different behavior patterns by changing the expression of a set of shared modules, rather than creating novel modules. In future work new modules might be discovered relative to landmarks other than the home. In larger environments, rodents are known to identify landmarks for navigation (Wallace et al., 2008), which might serve as anchors for the expression of some modules during foraging in the wild.

Our data indicate that complex economic patterns are comprised of discrete modules. Obesity, addiction, depression and anxiety disorders and others can be framed in terms of biological effects on economic patterns (Camerer, 2013; Hartley and Phelps, 2012; Monterosso et al., 2012; Rowland et al., 2008; Sharp et al., 2012b). We found multiple modules can link to a single primary outcome, like food consumption, raising the question of what functions these different modules serve and what internal states drive the expression of one module over another. Further, economic decisions during foraging involve tradeoffs (Stephens et al., 2007) and future studies could determine which modules have antagonistic relationships with each other and shape different tradeoffs. Defining the mechanisms involved in the form and expression of different modules could help uncover strategies to normalize maladaptive behavior patterns caused by genetic, epigenetic or environmental factors. Another goal involves developing “Ontological Categories” of behavioral modules based on their functions and analogs across species. Our study lays groundwork for this by developing eigenphenotypes as one approach for comparing modules across studies.

### Modular Changes to Behavior During Development

Much remains to be understood about how complex behavior patterns develop and change with age. This is an important area since most forms of mental illness involve early-life antecedents (Rutter et al., 2006) and arise at characteristic ages (Silbereis et al., 2016), and economic behavior patterns are necessarily linked to physiological and metabolic changes during development. Our data suggest that behavioral development for foraging involves creating and changing the expression of discrete modules. Currently, we do not know the functions of the modules or the mechanisms that underlie their development or expression at different ages. The genomics field has described developmental gene expression changes in the human, non-human primate and rodent brain (Bakken et al., 2016; Kang et al., 2011; Miller et al., 2014; Silbereis et al., 2016; Thompson et al., 2014), however, less is known about how specific mechanisms map on to specific behavioral changes. The majority of the cellular gene expression programs in the cortex of rodents and humans are highly conserved (Zeng et al., 2012), as are the trajectories of most neurodevelopmental gene expression programs in the developing brain (Bakken et al., 2016), and the temporal sequence of neurodevelopmental processes (Workman et al., 2013). Thus, the stereotyped trajectory of module development that we uncovered in mice could be rooted in conserved neurodevelopmental programs. Further, the expression of specific modules may help stage and study the progression of behavioral development in mouse models.

We found that parental effects on offspring foraging patterns unexpectedly emerge most strongly at ages after weaning. From an evolutionary perspective, parental effects are typically thought to be adaptive for parental fitness rather than offspring fitness (Haig, 2000; J Marshall and Uller, 2007) and the functions of parental effects on offspring behaviors after weaning are unclear. However, other adult behaviors have also been shown to be shaped by parental effects, including maternal behavior (Francis et al., 1999; Lefebvre et al., 1998; Li et al., 1999), circadian rhythm, feeding, reward, learning & memory and anxiety (Curley and Champagne, 2016; Perez et al., 2016). We speculate that parental effects on adult female foraging could function to influence foraging patterns in daughters during offspring rearing, contributing to adaptive, transgenerationally inherited economic patterns of behavior. Our study reveals candidate genes and pathways for further analysis.

Finally, by analyzing *Magel2* mutant mice we uncovered a genetic mechanism involved in regulating the expression of foraging modules and found that genes linked to human neuropsychiatric disorders can impact the expression of specific modules at specific ages. In humans, *MAGEL2* mutations cause PWS and autism (Schaaf et al., 2013), which involve major phenotypic changes across development (Cassidy et al., 2012; Lord et al., 2015). Further studies of *Magel2* mutants at other ages with our approach may help reveal the modular basis of behavioral changes due to mutations in this gene, and the mechanisms involved.

## STAR METHODS Animals

### Animals

Animal experiments were performed in accordance with protocols approved by the University of Utah Institutional Animal Care and Use Committee (#14-12012). C57BL/6J (B6) and CastEiJ mice were obtained from Jackson Laboratory. Mice were maintained on reverse 12 h light/dark cycle (lights off at 11:00) for foraging behavior experiments, and given water and food ad libitum (Harlan Tecklad rodent diet 2920X; Madison, WI.). All studies were performed on females. Cage bedding was Paperchip bedding (Shepherd Specialty Paper) and animals were housed with sex-matched littermates (2-5 mice per cage).

### Mouse breeding

Breeding pairs were housed together continuously for breeding. Maximum litter number was capped at 6 for each breeding pair and large litters were culled to a maximum of 6 pups. For P15 behavior testing, offspring were returned to the parental cage after testing and weaned at P21. For P20 behavior testing, mice were weaned after testing was completed. For P25 behavior testing, mice had been weaned at P21.

### Mouse Behavior Experiments

All B6, Cast, F1cb and F1bc mice were only tested once in the foraging assay and therefore mice at all ages were naïve to the test. Adult mice for all groups were at ~ 3 months of age when tested. *Magel2* mice were tested twice in the foraging assay, once at P20 and again as adults. For B6 females: N=20 adult, N=16 P25, N=15 P20 and N=16 P15 mice; For F1cb: N=17 adult, N=15 P25, N=15 P20 and N=15 P15 mice; For F1bc: 18 adult, 15 P25, 18 P20 and 15 P15; For Cast: N=15 P15 mice. For *Magel2* at P20, N=22 maternal wt (*Magel2*^m+’/p+^), N=28 maternal het (*Magel2*^m-/p+^), N=22 paternal wt (*Magel2*^m+/p+’^) and N=22 paternal het (*Magel2*^m+/p−^). For *Magel2* adults, N=18 maternal wt, N=23 maternal het, N=20 paternal wt and N=20 paternal het. Since each litter yielded ~ 3 females, each group involved testing pups derived from 5-9 different litters and at least five different breeding pairs. A detailed protocol for the behavior analysis is provided in the Supplemental Information.

### RNA Isolation and RNA-Seq

The DRN dissections include portions of the ventral periaqueductal gray and the RNA isolation, RNA-Seq and imprinting studies were performed as previously described (Bonthuis et al., 2015; Huang et al., 2017). In brief, RNA was extracted and DNase treated using the RNAeasy Micro Kit (QIAGEN). RNA was pooled from ~6 daughters from different litters to provide 3 μg of total RNA for each biological replicate. Samples were prepared for RNA-seq using the TruSeq RNA sample preparation kit v2 (RS-122-2001, Illumina). Single-end 59-bp sequencing of the libraries was performed using the Illumina HiSeq 2500.

### Statistical Analysis of Behavior

Statistical testing was performed in R and the code and sample datasets are provided in the detailed protocol file in the Supplemental Information.

### Statistical Analysis of RNASeq Data

Statistical testing of gene expression differences in the RNASeq data was performed using the edgeR package (Robinson et al., 2010).

## SUPPLEMENTAL INFORMATION

Supplemental Information includes 2 supplementary movies, 8 supplementary figures and 6 tables, supplemental data and a detailed protocol for the foraging assay with code and 3D printing files.

## AUTHOR CONTRIBUTIONS

Conceptualization: C.N.S.H and C.G., Methodology: C.N.S.H., E.W., P.T.F. and C.G.; Investigation: C.N.S.H and A.N.R; Software: C.G., C.N.S.H., E.W.; Formal Analysis: C.N.S.H, E.W., P.T.F., E.F., C.G.; Visualization: C.N.S.H., E.W. and C.G.; Writing: C.N.S.H., E.F. and C.G.; Supervision: P.T.F. and C.G.; Funding: C.G.

## ACKNOWLEDGEMENTS

We wish to thank Drs. Matt Wachowiak, Denise Dearing, Jan Christian, Jared Rutter, Carl Thummel, Megan Williams, Adam Douglass, Emilie Rissman, Gaby Maimon, Margaret McCarthy, Monica Vetter, Rich Dorsky and the Gregg lab for comments on earlier versions of the manuscript. C.N.S.H. was supported by a Swiss National Science Foundation Postdoctoral Fellowship and the Gottfried & Julia Bangerter-Rhyner Stiftung. E.W. and P.T.F are supported by NSF grant IIS-1251049. C.G. is a New York Stem Cell Foundation Robertson-Neuroscience Investigator and supported by NIMH R01 MH109577.

---

**Figure 1-S1.**
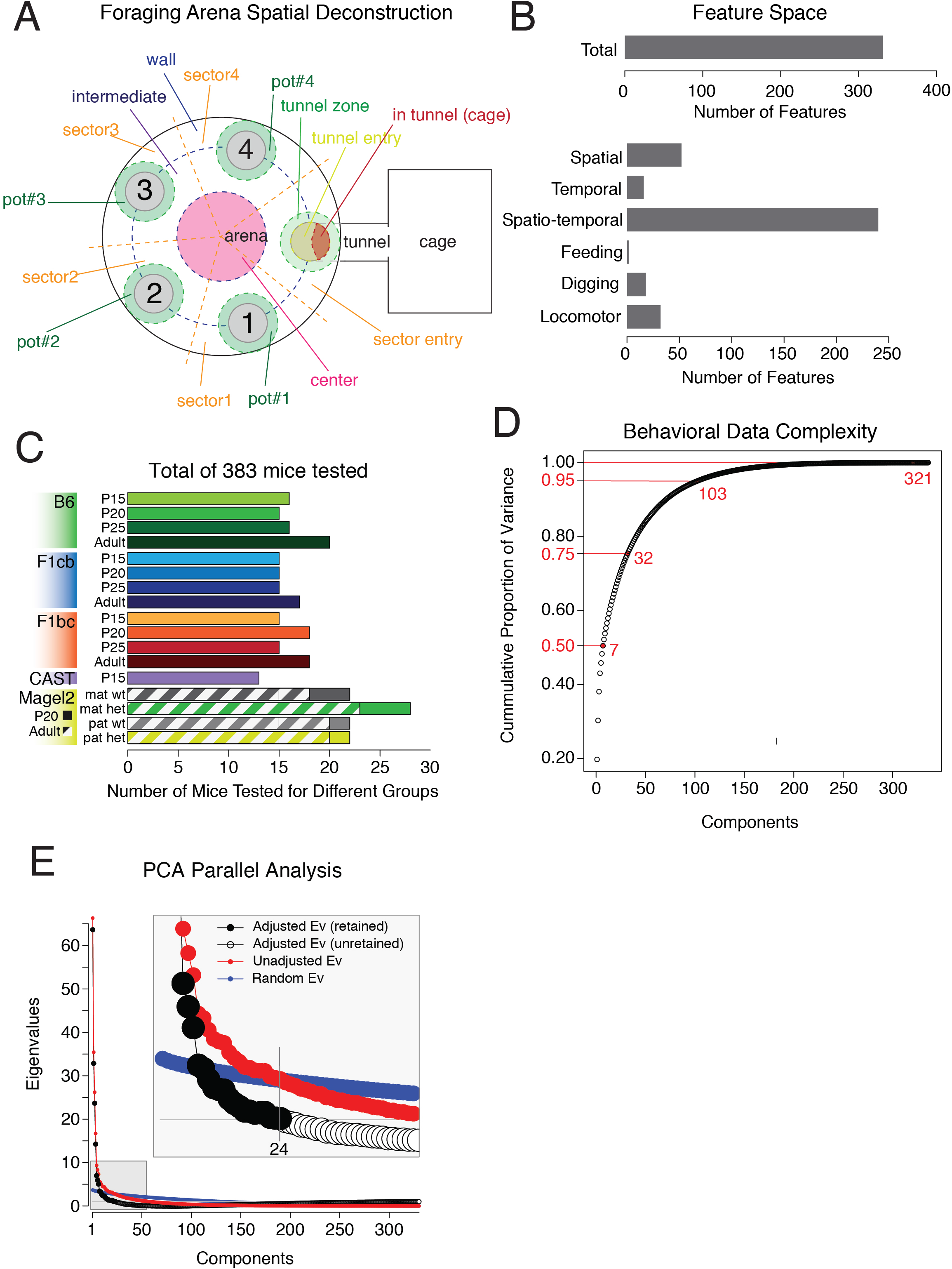
Identification of a high dimensional feature space for the analysis of complex foraging patterns. **(A)** Schematic overview of the different zones of the arena for systematically deconstructing spatiotemporal features of foraging. **(B)** The bar chart shows the total feature space (338) and a break down of numbers of different types of features defined according to different types of measures (see Table S? and Supplementary Information). **(C)** The bar chart summarizes the number of female mice profiled for each age and genotype in our study (C). **(D-E)** A PCA analysis of the feature data for all 383 mice reveals high dimensionality for the data. The dot plot in (D) shows the cumulative proportion of the variance that is explained by different numbers of principal components. The plot in (E) shows the results of a parallel analysis, in which 24 principal components are retained to explain the variance feature data. Ev, eigenvalue.

**Figure 1-S2.**
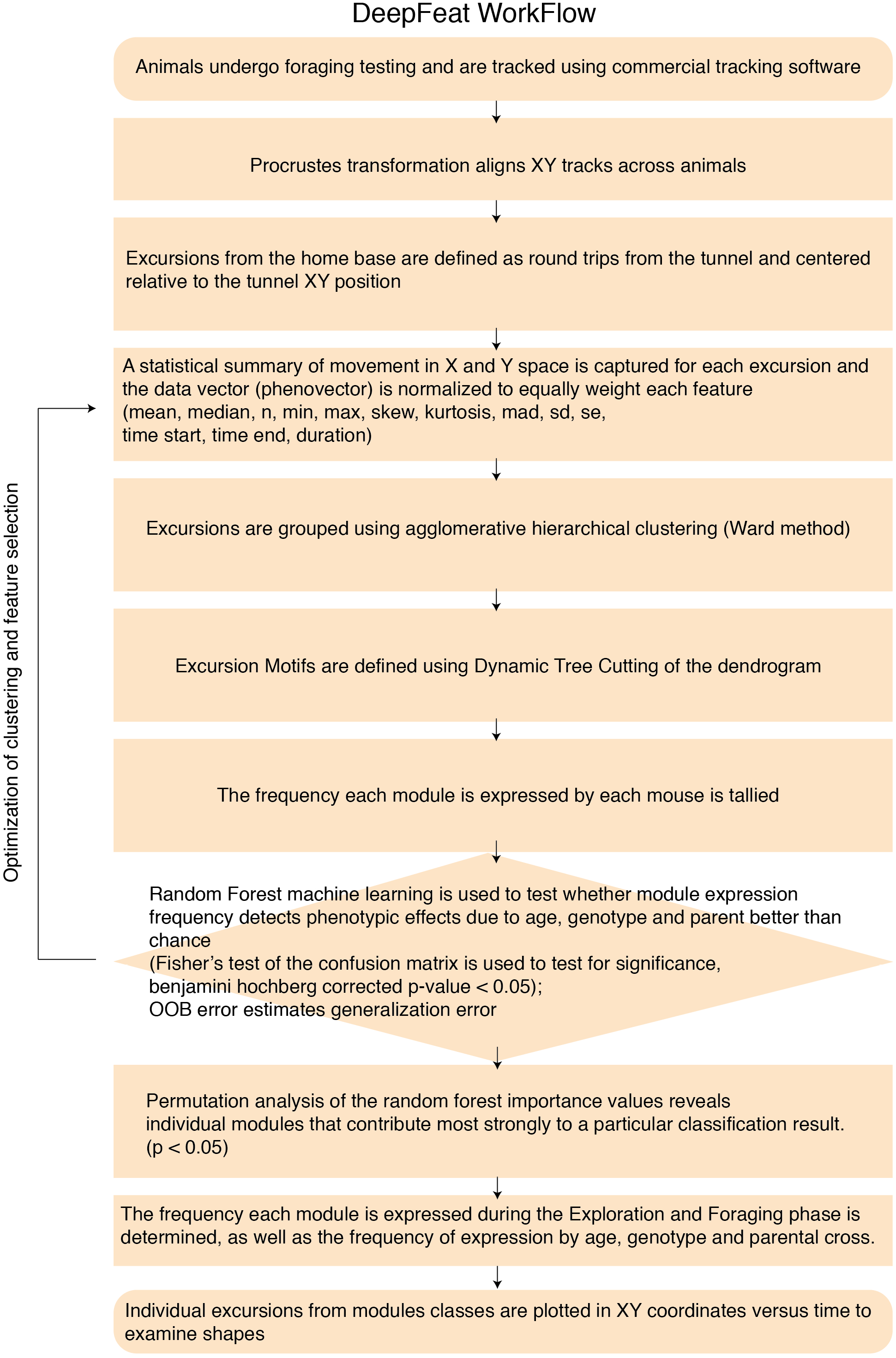
Summary of the workflow for DeepFeat analysis of behavioral modules. The graphic shows the different steps involved in DeepFeat. Custom scripts were generated to perform each step of the analysis.

**Figure 2-S1.**
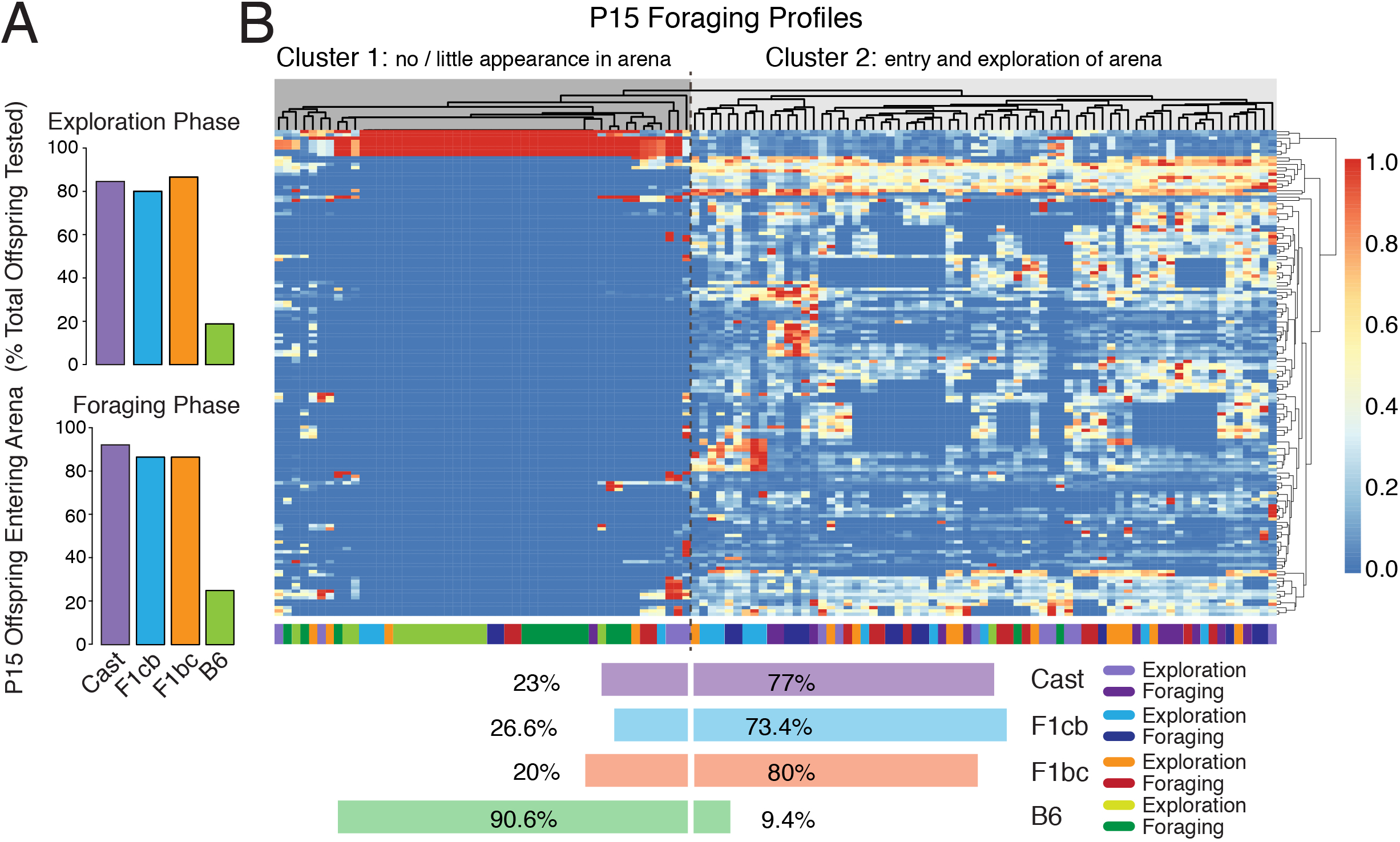
Foraging is immature in P15 mice, but early stage patterns are detectable from many features for P15 Cast, F1cb and F1bc offspring. **(A)** Bar plots show the percentage of P15 juveniles that appear in the foraging arena in the Exploration phase and Foraging phase (N=15-20 per cross). Few B6 offspring (20%) enter the arena, but most Cast, F1cb and F1bc (~80%) offspring enter. **(B)** Heat map and clustering results for the profiles from all Exploration and Foraging trials for B6 (green), Cast (purple), F1cb (blue) and F1bc (red) P15 animals. Columns are single trials. Rows are behavioral measures. Values are normalized to maximum value in each row. Colored bars underneath the heatmap indicate the cross (B6, Cast, F1cb, F1bc) and phase for each trial (column). Horizontal bars indicate the percentage of trials from each cross that cluster into Cluster 1 (no/little appearance) or Cluster 2 (data captured from foraging trials). The results in Cluster 2 show that data for many features was successfully obtained for P15 Cast, F1cb and F1bc offspring.

**Figure 2-S2.**
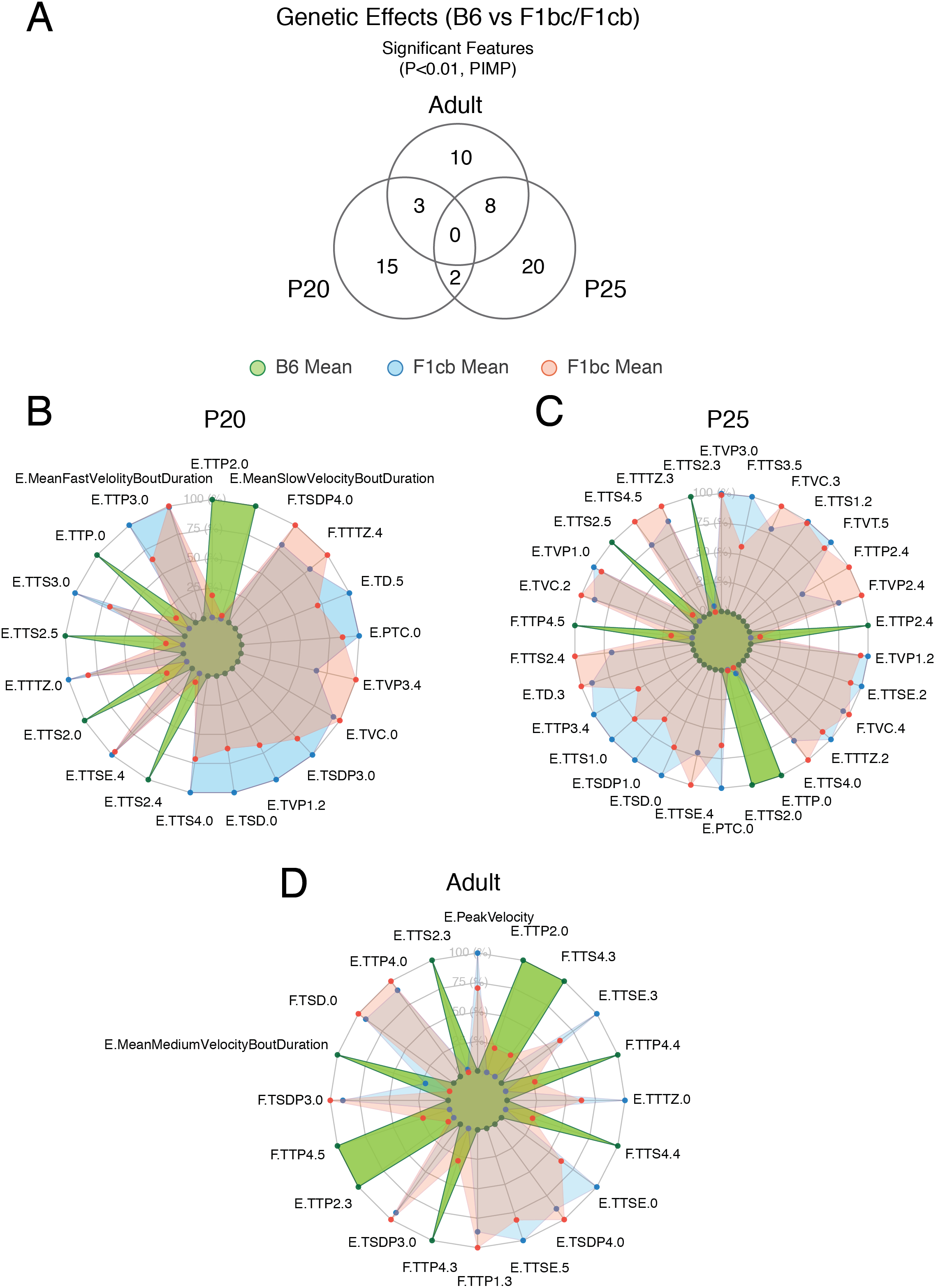
Genetic effects in B6 versus F1 hybrid mice shape foraging patterns in an age-dependent manner. **(A)** A venn diagram of the features with significant importance values in the random forest classifiers detecting B6 from F1 hybrid offspring at P20, P25 and adult. No feature is significant across all ages and most features are age-specific. **(B-D)** The radar plots depict the data for a subset of the top ranked features (PIMP P<0.01) in the classifier detecting B6 offspring from F1cb and F1bc offspring at P20 (B), P25 (C) and for adults (D). The data points for each behavioral feature are the mean values from B6 (green), F1cb (blue) and F1bc (orange) mice, presented as percentages of the relative maximum (100%) and minimum (0%) values for each feature. Feature codes are: [Phase.Feature.TimeBin]. F, Foraging phase; E, Exploration phase. See Table S1 for feature definitions.

**Figure 4-S1.**
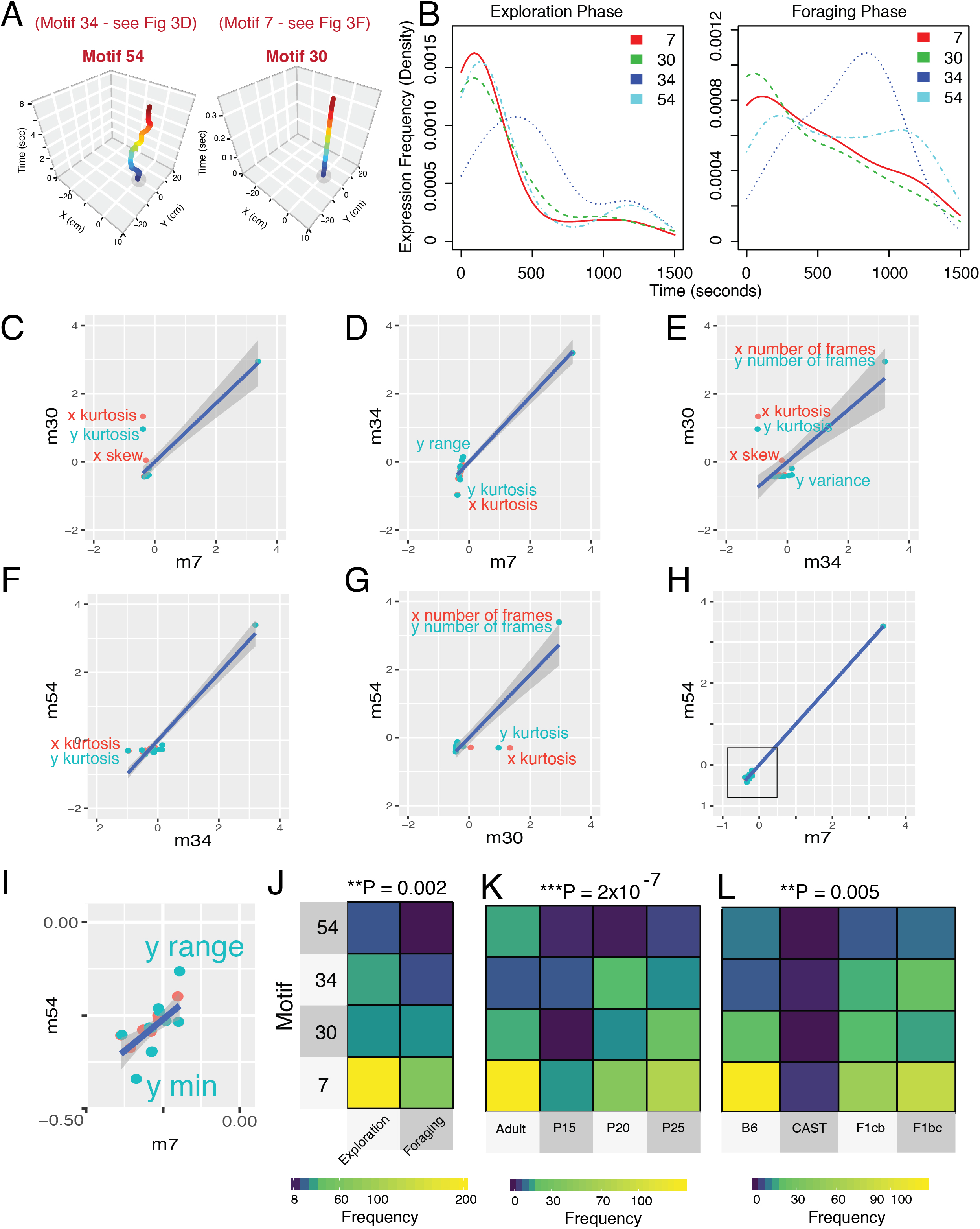
Modules involving brief behavioral sequences capture important phenotypic information. **(A)** Four short modules were analyzed in depth. Representative traces are shown for two of the short modules, including Module 54 and Module 30. Traces for Module 34 and Module 7 are presented in other main figures. The short module subtypes show differences in temporal length and/or XY movement architecture. **(B)** Density plots of the expression pattern (excursion start times) for all excursions classified as Modules 7, 30, 34 or 54. The temporal expression pattern is shown across the entire 25 minute Exploration and Foraging phases. The data shows that Module 34 exhibits a distinct temporal expression profile in the Foraging phase compared to the other short modules. **(C-I)** Plots compare descriptive statistics calculated from the XY movement patterns for excursions falling under the same module class. Data for each module (m) is plotted in a comparison with a different short module (7, 30, 34, 54). The results show specific differences and outlying measures relative to a regression line indicating the pattern for a linear relationship. The data in (I) is a zoom of the data in the black box in (H). Prominent outlying descriptive statistics are labeled in each contrast. **(J-L)** The heatmaps show the frequency each short module is expressed according to phase (J), age (K) and genotype (L). Different modules show different expression patterns. A Chi-Square test of independence determined that the short module expression frequencies are significantly dependent on the biologically meaningful factors of phase, age and genotype (P values shown on top).

**Figure 5-S1.**
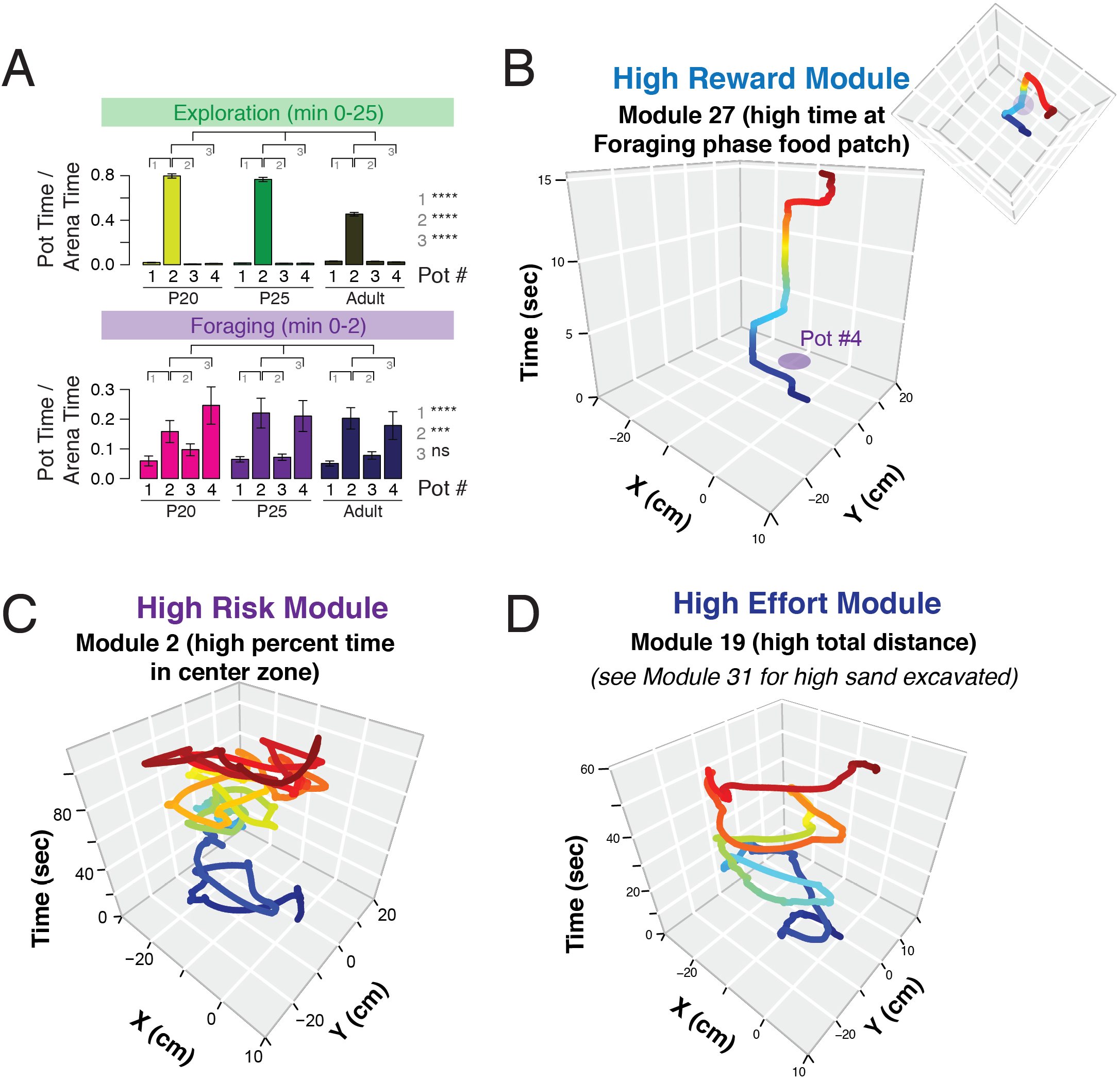
Top modules linked to reward, risk and effort responses during foraging. **(A)**Bar charts show the time spent at each pot (1, 2, 3 and 4) during the entire Exploration phase and the first two minutes of the Foraging phase. The time at each pot is normalized to the total time an animal spends in the foraging arena. The data show that P20, P25 and adult mice find the food in Pot#2 in the Exploration phase and retain a perseverative memory of that food’s location in the Foraging phase. Significance relationships are shown in the legend. One way ANOVA and Tukey posttest. N=20-25 B6 mice. Error bars show SEM. The data show memory and perseveration responses, providing the basis for the identification of putative memory and perseveration modules. **(B-D)** Representative traces of the top modules linked to total food consumption (B), percent time in the center zone (C) and total distance traveled (D). The inset rotation of Module 27 shows overlap with the Foraging phase food patch in Pot #4 (B). Module 2 is characterized by extensive time in the center zone of the arena (see Fig. 5-S1 for location of center zone). Module 19 involves long, wide looping movements.

**Figure 7-S1.**
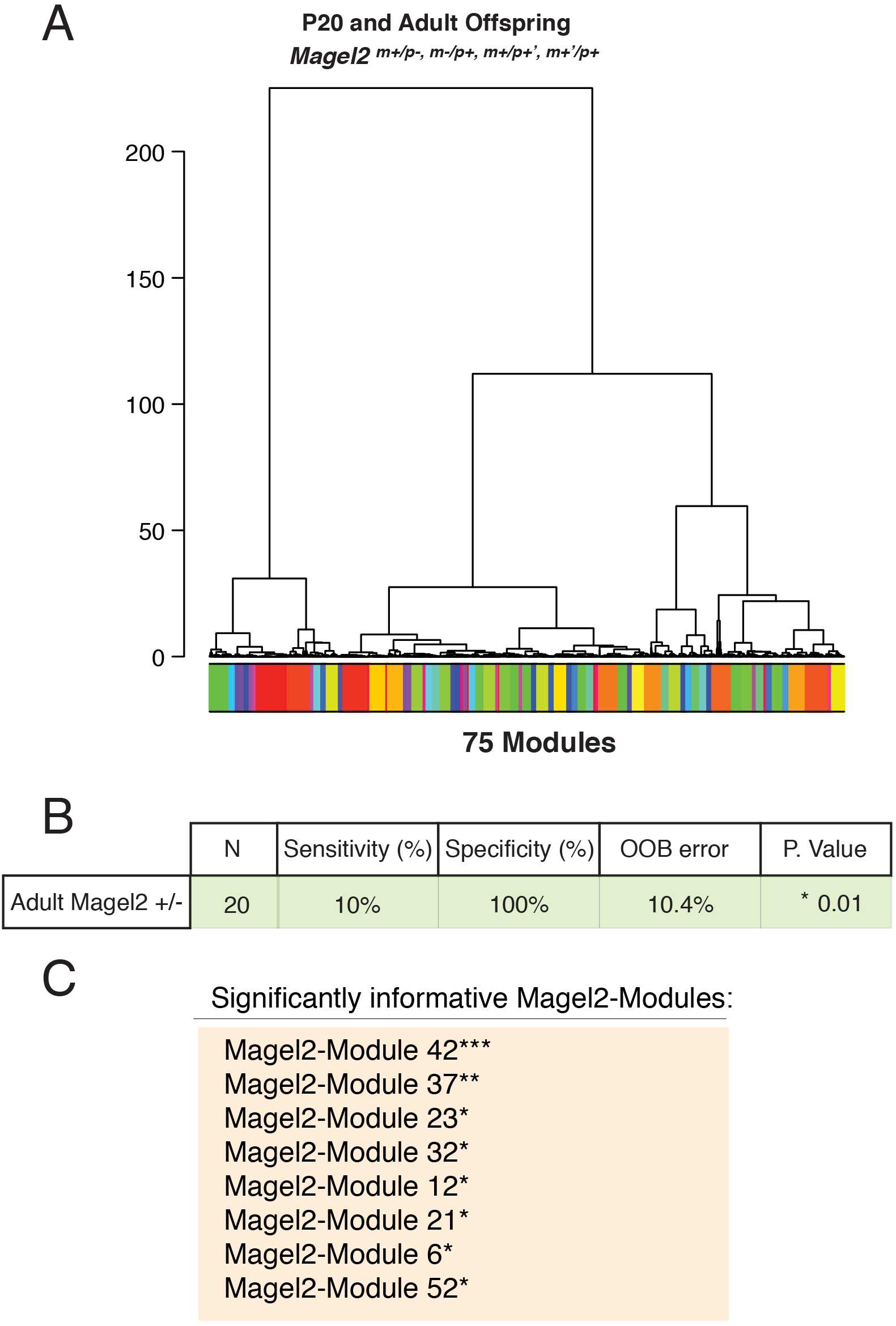
Identification of behavioral modules expressed by P20 and adult *Magel2* mutant and control mice. **(A)** Dendrogram of the clustering results to define modules from home base excursions. DeepFeat identified 75 modules from 5302 home base excursions performed by 60 mice in our cohort of P20 and adult *Magel2* mutants and controls. Distinct clusters (modules) are indicated by color. **(B)** The table shows the results of the random forest classification of adult *Magel2^+/−^* mutants based on module expression frequency data. A two-tailed Fisher’s test on the resulting confusion matrix showed statistically significant detection of adult *Magel2* mutants (p-value shown). OOB, out-of-bag error. **(C)** The table shows the identity of the nine significantly informative Magel2-Modules for the adult *Magel2* mutant phenotype found from permuted importance analysis. ***P<0.0001, **P<0.01, *P<0.05

**Figure 7-S2.**
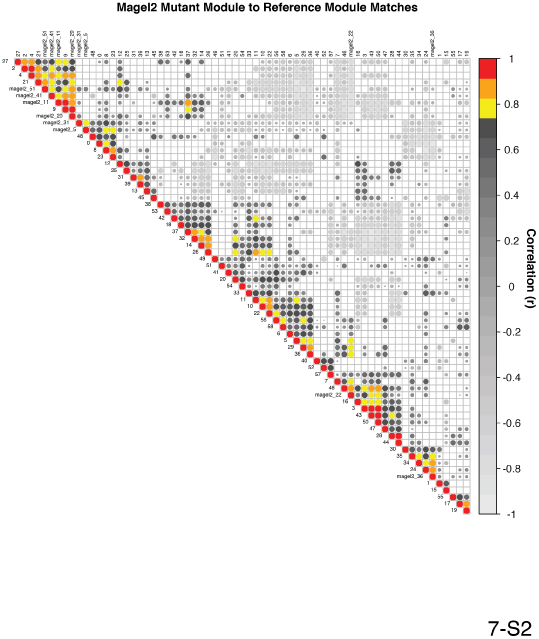
Eigenphenotype correlations reveal matches between Magel2-Modules and Reference Modules derived from Cast, B6, F1cb and F1bc mice. The plot shows the upper triangle of the correlation matrix matching nine Magel2-Module query modules to all 59 core reference modules. Top hits are colored according to the legend and the best reference module matches for each Magel2-Module are shown in Fig. 7H. Only statistically significant matches are shown in the plot after adjusting for multiple testing. Holmes adjusted p-value < 0.05

## SUPPLEMENTAL MOVIE LEGENDS

### Movie S1: Exploration trial

Movie showing short sequences of a representative adult Exploration phase trial. The adult B6 female appears in the arena, retreats to the home cage and returns several times to the arena. The mouse explores various features and regions of the arena environment and finally finds the food in Pot #2 and starts to eat. This phase examines how mice behave in a novel foraging environment and discover and consume food. We deconstruct the behavior patterns in this phase into 152 different measures.

### Movie S2: Foraging trial

Movie showing short sequences of a representative adult Foraging phase trial. The adult B6 female appears in the arena and visits the previous food pot (Pot #2), digs and searches for food in Pot #2. The mouse eventually gives up searching Pot #2, searches and finds the hidden food in the new location (Pot #4). This phase examines how mice behave in a more familiar foraging environment, adapt to changes in the environment and discover the novel food source. We deconstruct the behavior patterns in this phase into 152 different measures.

## SUPPLEMENTAL TABLES

**Supplemental Table 1. Summary of all behavioral and locomotor features measured.** For Figures 1-7. This table details the features measured in the foraging assay. Each feature is measured in the Exploration phase and the Foraging phase and several measures are analyzed at different time bins (1, 2, 3, 4 and 5) and/or across the entire phase (0). Each measure is assigned a unique ID with the following structure [Phase.Feature.TimeBin].Tab#1 Behavior Features: phase, Exploration or Foraging phase; measure, description of each behavioral feature; code, unique ID for each behavioral measure; unit; non-normalized data, x indicates if a measure is included in the non-normalized data set; normalized data, x indicates if a measure is included in the normalized data set; normalization, values indicate how each measure was normalized for the normalized data set.Tab#2 Locomotor Features: The locomotor features captured in the assay are indicated with descriptive IDs.

**Supplemental Table 2. Mouse phenome project detected phenotypic differences between Cast and B6 female mice that could contribute to putative parental effects on offspring phenotypes.**For Figure 2A. The table shows phenotypic differences between Cast and B6 female mice uncovered by the Mouse Phenome Project. Significant differences in female reproductive behaviors were identified, as shown. These phenotypic differences are diverse and support the rationale for investigating parental effects on offspring in F1cb and F1bc mice.

**Supplemental Table 3. Raw data for Cast, B6, F1cb and F1bc foraging features and motifs.**For Figures 2-5. The raw data for features as defined in Table S1 and for motif expression counts identified by DeepFeat.

**Supplemental Table 4. Significant differentially expressed genes identified by RNASeq analysis of F1cb versus F1bc female mice at P5, P15 and adulthood for the DRN region.**For Figure 6. The results show the output from edgeR for the analysis of differential gene expression due to parental effects at different ages. Separate tabs indicate the data for P5, P15 and adult female mice. CPM, counts per million reads; FDR, false discovery rate.

**Supplemental Table 5. DAVID gene ontology and pathway enrichment results for genes differentially expressed in the DRN between F1cb and F1bc mice at P5 and Adult.**For Figure 6. The ontology enrichment analysis was performed on differentially expressed genes defined at a 10% FDR threshold relative to a background of all genes expressed in the DRN. Separate tabs are provided for the P5 and adult DRN results.

**Supplemental Table 6. Raw data for Magel2 mouse foraging features and motifs.**For Figure 7. The raw data for features as defined in Table S1 and for motif expression counts identified by DeepFeat.

## SUPPLEMENTAL INFORMATION

### Rationale for the Foraging Assay Design and Feature Selections

#### Foraging Assay Design

Our primary goal in the design of the assay for the foraging screen is to create a broad, ethologically-relevant screening platform in which diverse phenotypic patterns can potentially manifest due to effects from a wide variety of different neural systems. In our design, we attempt to incorporate important elements from behavioral ecology and lab-based approaches to the analysis of foraging. To investigate rodent foraging in the wild, behavioral ecologists frequently monitor trays of sand that contain seeds (Brown, 1988b). These food patches are placed strategically in different regions of the environment to test different hypotheses about rodent foraging patterns and to measure giving up densities. The giving up density is the point at which rodents abandon a food patch because the density of seeds in the sand is too low, and therefore the animal has made a decision about the costs of continued foraging in that patch versus the rewards (Brown, 1988a; Brown and Kotler, 1994a; 1994b). We incorporate this idea into our assay by designing the food patches in the foraging arena as pots of sand with or without seeds. Thus, mice are challenged to extract seeds from the sand and apparently make decisions regarding the costs versus the rewards (see below). Our design also incorporates features from several standard lab tests (Wahlsten, 2010), including the open field test (thigmotaxis/activity), marble burying/digging (repetitive/perseverative behaviors), Barnes maze (spatial navigation/memory), Morris water maze (spatial memory/reversal learning), the buried food test (sensory) and basic food consumption analysis (hunger/satiety), as we describe further below. From relatively free and spontaneous foraging, these different components of the design create an opportunity for diverse behavioral patterns to manifest in mice, as we show in our study.

We wish to minimize the handling of the mice and allow them free access to the home base cage during the foraging trials. Therefore, we designed the home base cages with a tunnel and door (see Protocol – Arena Construction). The foraging arena is also designed with a tunnel, which leads into the arena, and during a trial, the home cage tunnel plugs into the arena tunnel, the door is opened and the mouse can now freely leave the cage to access the foraging arena through the tunnel, and vice versa. An important point to note is that in a typical lab-based behavior test, the mouse is captured, lifted and placed into a behavioral testing apparatus. However, in the wild, animals freely and spontaneously move between their home base and the outside environment and the home base is an important safe shelter that can shape exploratory patterns (Eilam and Golani, 1989).

We hypothesize that genetic mechanisms in mice shape many different foraging behavior patterns relative to the home base and therefore this is an important element of our design. Some known genetically driven behaviors associated with the home include territorial behaviors, burrowing and nest construction (Sauce et al., 2012; Weber et al., 2013), but other genetically-driven behavioral patterns likely exist. We attempt to capture elements of home base related behaviors during foraging in our screen by deconstructing spatiotemporal features related to the home base. For example, each round trip from the home base into the arena and back is considered a single excursion (Drai et al., 2000) and many discrete features of these excursions from the home base are analyzed in our assay. We capture data for the length of excursions from the home base at five different time windows during the assay, time spent in close proximity to the home base, the number/frequency of excursions, the length/frequency of fast velocity, medium velocity and slow velocity bouts of movement in each excursion from the home base, the duration of time away from the home base, excursion patterns from the home base during the naïve Exploration phase versus the more familiar and goal-directed Foraging phase, preferences for exploring potential food patches closer to the home base and the latency to enter the arena from the home base. These are some examples of patterns and features that could be shaped by genetic mechanisms that influence home base related behaviors.

Optimal foraging theory makes predictions about the strategies animals should use during foraging to maximize rewards and reduce costs while seeking out and exploiting food patches (Charnov, 1976; Pulliam and Charnov, 1977). Seeking out new food patches is risky, since it is unclear whether new resources will be uncovered during a search and the effort can result in energy expenditure and predation costs. Thus, familiar foraging environments where known resources exist are valuable, and an animal faces the ongoing challenge of deciding when to give up on a known food source and take the risk of seeking out a new resource. Many neural systems are engaged in these aspects of foraging, including neural systems for exploring, navigating and detecting resources, learning the location of resources in the environment, remembering the location of resources and giving up on food patches that are exhausted (Stephens et al., 2007). Our assay is designed to capture some patterns related to these important aspects of foraging, as follows.

First, mice search pots of sand in order to discover and obtain seeds for food. This aspect of the design is ethologically relevant and innate for mice, but it also demands effort, since the animals dig through the sand to explore, discover, extract and consume seeds. Secondly, by incorporating a naïve Exploration phase with an obvious food patch and then a relatively more goal-directed and familiar Foraging phase, in which the food patch is switched and hidden, we expect that the assay can reveal a variety of different phenotypic effects and patterns. For example, we expect to uncover patterns related to: (1) the discovery of a food patch (Pot#2) in a new environment (Exploration phase), (2) the subsequent return to, investigation of and then abandonment of a former food source (Pot#2) in a now familiar environment (Foraging phase), and (3) the discovery and exploitation of a new hidden food patch (Pot#4) in a familiar environment (Foraging phase). In the current design, mice are provided with a 4 hour break between the Exploration and Foraging phases, which challenges the formation and maintenance of memories of the food location beyond short-term working memory, but allows for several animals (~6-8 animals) to be tested within a single day, which scales to ~48 animals per week for screening (see Protocol). Each testing phase is 25 minutes long, which is designed to allow for some adaptation and the expression of different early, mid and late stage behavior patterns, which we show occurs in the main text.

In addition to using the safety of the home base, animals have innate behaviors to find food and avoid predation during foraging that influence how they navigate and behave in exposed spaces versus less exposed spaces in the environment. One example of this dimension of behavior is thigmotaxis, which involves a natural preference for walls in an otherwise exposed area and occurs in mice (Prut and Belzung, 2003) and humans (Walz et al., 2016). Our arena is designed with an exposed center region and wall zones to capture behavioral features related to this dimension of foraging. Further, the pots are positioned so that animals have the option of approaching, exploring, digging and feeding at the pots while their body is positioned toward the wall or out toward the exposed center. To capture data for features potentially shaped by thigmotaxis, we break the arena into concentric circles (Fig. S1A) that capture visits and the duration of time in (1) the most exposed center region, (2) in an intermediate zone where the animal can interact with the pots with its body positioned toward the center, and (3) a wall zone where the animal positions its body close to the wall. By analyzing behavioral patterns in these different zones, we potentially learn about some strategies the animal is using to reduce risks of exposure during the Exploration and Foraging phases of the assay.

Finally, an important point regarding the design is that rodents explore and forage instinctively, and do not require food deprivation to motivate these behaviors (Collier et al., 1972; Kavanau, 1967; Moran, 1975; Pierre et al., 2001); however an increased motivational state enhances learning (Makowiecki et al., 2012), which is a component of our assay. Therefore, we food deprived the mice prior to the assay such that the animals experience the same degree of body weight loss (~10% of starting body weight), setting a common baseline for comparison across ages (see Supplemental Experimental Procedures). A second point to note is that young animals (P15 and P20) are maternally deprived since they are removed from the mother during the assay, which could shape their behavior patterns. However, maternal attachment changes and the anxiety that arises due to maternal separation declines postnatally and is a natural part of the development of independent foraging behavior (Galef, 1981).

#### Heuristic Feature Selection for Pattern Detection from Machine Learning

The approach we took to selecting features for analysis in our assay is intended to achieve three main goals. First, we wish to uncover novel behavior patterns that would not have been predicted *a priori.* Second, we wish to be able to interpret and have at least some understanding of the observed patterns, the features involved, and their potential relevance to the principle objective of foraging – the discovery and consumption of food through strategies that reduce effort and predation costs. Some machine-vision based approaches can potentially uncover complex patterns that are challenging to interpret, and we sought to reduce this in our initial screen, though it remains an area for future development. Finally, we wish to be able to study and compare behavior patterns across ages, and therefore between mice with different sizes and shapes. For this application, the features and detected patterns need to be comparable and generalizable across ages, which can also be challenging for some machine-vision based strategies (Kabra et al., 2012; Wiltschko et al., 2015).

To meet these objectives, we first systematically deconstruct behavior patterns in the assay into progressively more restricted spatiotemporal features (Fig. S1A). Spatial features are deconstructed first by breaking the arena into pie shaped sectors that encompass each major element of the arena (the pots and tunnel entry zone). Next, we define regions immediately surrounding each major element in the arena (pot zones and the tunnel zone). The tunnel zone is further subdivided into an entry zone, where the mouse may peak into the arena from the tunnel and a tunnel zone, where it is in the arena, but near the tunnel. Finally, we break the arena into domains according to two concentric circles, one small circle encompassing the exposed center region and a second larger circle that divides the pots and tunnel zone in half, with one half proximal to the wall and the other half facing the center. This approach systematically divides the arena into a large number of increasingly restricted zones with interpretable significance.

Temporal patterns are derived by collecting data for visits and time spent at different zones in the arena in five 5-minute time bins for each phase and in aggregate over the entire phase. In our main study, we found that 5-minute time bins perform well for phenotype detection and are relatively easily interpreted as early, mid and late stage phenotypic effects. Nonetheless, it is possible that new phenotypic information could potentially be uncovered by further atomizing behaviors with finer temporal resolution. Additional, temporal dimensions of behavior patterns are revealed from latency measures, as well. As an illustration of how these different temporal dimensions might find different patterns, a mouse may obtain a lot of food from Pot #2 during the Exploration phase by simply stopping there and eating, which could be reflected in a long duration of time at Pot #2. Alternatively, the mouse could obtain a lot of food by darting in and out quickly from the food pot many times and performing many short visits. For this sort of potential effect, examining both visits and durations at different areas in the arena at different times during the assay can potentially reveal important patterns. Similarly, mice may initially spend a lot of time near the tunnel to the home base during the Exploration phase, but gradually become comfortable exploring pots further away from the home base cage (Pots #2 and #3) at later stages, performing longer and longer excursions into the arena. They may sprint through an exposed region, like the center, to reach an objective at early stages, like a pot of interest, or they may slowly creep around the arena, cautiously and systematically exploring. Overall, we don’t know which features or patterns might be shaped by particular genomic elements or what patterns might emerge at different times. Therefore, we systematically capture visit, duration and latency data for all of the different regions defined in the arena.

Locomotor features and patterns are major component of our analysis of foraging phenotypes with our screen. Locomotor patterns can potentially reflect diverse phenotypic effects that could be related to anxiety & fear levels, activity, motivational state, motor patterns, anatomical phenotypes and many other effects (Drai et al., 2000; 2001; Fonio et al., 2009; Kafkafi et al., 2003). Indeed, human studies have found important and unexpected relationships between motor patterns and various psychiatric symptoms (Peall et al., 2017). In the current version of our screen, we analyze the length, frequency and duration of slow, medium and fast bouts of velocity at different stages of the testing periods. We also classify Exploration and Foraging phase excursion patterns into major subclasses based on the elements of the arena that are visited, the distance traveled, the duration of the excursion, and the frequency/duration of the different velocity bouts. By deeply analyzing these different features of locomotion we expect to be able to uncover diverse phenotypic effects that manifest in this dimension of foraging behavior.

Finally, food consumption and digging behaviors are important features of foraging that can be shaped by a variety of phenotypic effects. For example, while hunger can increase food consumption, stress & fear can potentially reduce the amount of food consumed (Brown and Kotler, 2007). Additionally, digging is a normal part of mouse exploratory and foraging, but could indicate abnormal activity, repetitive or perseverative behaviors in our assay. By analyzing visits, time spent and the mount of digging in Pots #1 and #3, which don’t contain food, we may learn about the animal’s strategies for exploring other potential patches even when an abundant and obvious resource has been uncovered (eg. Pot#2 in the Exploration phase). Thus, measuring the total sand dug from each pot and the total food consumed during each phase provides us with important insights into the behavioral patterns expressed and their impact on the central goal of the behavior – finding and consuming food.

Our discussion above explains the rationale for the assay design and the features we detect. While there are many dimensions of natural foraging that we cannot capture, our screen nonetheless examines a variety of different behavioral and locomotor features to detect diverse patterns that we would not have predicted *a priori*. In total, we obtain 336 features from the assay and each is assigned a unique ID (Table S1). We do not claim that the features we defined are the only important features that could be extracted from our assay and we expect that future studies will continue to resolve new and important features of foraging in our screen. However, in the main text, we provide evidence that our approach robustly detects a variety of different phenotypic effects at different ages. For some phenotypes, some features may be redundant, but they are not necessarily redundant and might reveal important phenotypic information in other mouse models not tested in the current study. Importantly, random forest ensemble learning, which we use to detect distinguishing behavioral patterns, is not confounded by correlated features or high dimensional data, and therefore the addition of more spatiotemporal and locomotor features simply expands the landscape for pattern discovery.

### Known Phenotypic Differences between Cast and B6 Mice Could Potentially Contribute to Parental Effects on Offspring Phenotypes

By analyzing data in the Mouse Phenome Database (Grubb et al., 2013), we found that Cast females have a significantly lower Parenting Index (ratio of weaned to born pups) compared to B6 females, indicating differences in maternal behavior/physiology that impact offspring postnatal survival (Table S2). In addition, Cast and B6 females differ significantly in terms of litter size, fecundity, body size, hormone levels, metabolic traits, activity, body composition, immune cells and cardiovascular traits (Table S2). Differences in these traits can influence offspring development (Asvold et al., 2011; Boersma et al., 2014; Lim et al., 2014; Moog et al., 2015; Schlotz and Phillips, 2009; Westberg et al., 2016). Beyond behavioral/physiological differences, cis-regulatory differences between Cast and B6 mice cause differences in the expression levels of some imprinted genes in F1cb versus F1bc offspring (Bonthuis et al., 2015), and mitochondrial genes are Cast-derived in F1cb mice, but B6 derived in F1bc mice. Thus, F1cb and F1bc offspring have various potential differences in environmental and genetic parental effects that could shape offspring behavioral phenotypes in our screen.

## SUPPLEMENTAL EXPERIMENTAL PROCEDURES

The initial prototype foraging assay design and protocol, as performed in this study, is provided below. In a separate supplemental Protocol file, we provide a detailed protocol for an updated arena design, assay methodology and computer code for the data analysis.

### Foraging Assay Protocol

In preparation for the foraging assay, mice were first habituated with sand (Jurassic play sand, Jurassic Sand) and seeds (Whole millet, Living Whole Foods) for two days in their home cage. On day one, seeds are spread on top of sand in the bottom of a petri dish and the dish is placed on the bedding in the home cage for the mice and pups to explore. On day two, seeds are covered with sand in the bottom of the petri dish in the home cage for the mice and pups to dig in and explore. To motivate animals to feed, mice were food deprived prior to testing to achieve 8-10% weight loss at the time of testing. For P15, P20 and P25 mice, weight gain normally occurring during the period of food deprivation was taken into account to achieve the intended weight loss. We selected this weight loss target after several pilot studies with the goal of achieving some consistency in the motivational states of the animals at different ages and not compromising health or activity. To achieve the intended weight loss and motivational state, adult mice were food deprived for 24 hours, P25 mice were food deprived for 17 hours, P20 mice were food deprived for 17 hours and housed with the mother, P15 mice were food deprived for 4 hours prior to testing in the absence of the mother. For food deprivation, mice are moved into a fresh cage with some soiled bedding to ensure familiar smell, but no leftover food in the cage. Water is available ad libitum at all times except when mice are in testing cage (2 × 1 hour).

Mice are housed in a room with an 11:00 – 23:00 dark cycle, so that testing is performed during the dark cycle. For testing, mice are moved into the behavior room prior to the start of testing for at least 1 hour for habituation to the new room. All testing is performed in the dark and video recording is done using infrared illumination and all manual procedures are done in the dark using a headlamp with red light. The mouse to be tested, and their home cage soiled bedding, are moved to the testing-cage attached to the arena 30 minutes prior to testing and allowed to habituate. At the start of testing, the testing-cage is attached to the arena via the tunnel, the mouse now has access to the arena and video recording starts for the Exploration phase. Mouse behavior is recorded continuously during the 30 min Exploration phase trial under infrared lights. Noldus Ethovision software v8 and v9 were used for video tracking. After completing the Exploration phase, the mouse is transferred to a holding cage with water but no food until the Foraging phase four hours later. For the Foraging phase, mice are again placed in the testing cage and habituated for 30 min. The testing cage is then gently attached to the tunnel and access to the arena is possible and the Foraging trial begins. Video recording of the Foraging phase is performed for 30 minutes. After testing, mice are placed in a new cage with food and are returned to the mouse colony room. Importantly, between each Exploration and Foraging phase trial, the entire arena, including walls, platform, tunnel and steel pots, are wiped clean with 70% ethanol.

### Preparation of Sand and Seed Pots for the Foraging Assay

Three stainless steel pots (Resco, diameter 5.5cm, depth 4cm) were filled with 95g of sand. For the Exploration phase, 1 pot is filled with 80g of sand covered with 2.5g of seeds. On top of seeds, a layer of 12g of sand is added to cover seeds. This sand is then covered with 0.5g of seeds. This pot is placed in position 2 in the arena. For the Foraging phase, 1 pot is filled with 80g of sand, 3g of seeds on top of sand and additional 12g of sand to cover all seeds. This pot is placed in position 4 in the arena. All pots are weighed before and after the trial to measure the sand displaced from each pot. Remaining seeds and hulls left in the pot and on the platform are measured after each Exploration or Foraging trial to determine the amount of seeds consumed by the mouse during the trial. Used sand is collected after every trial and set aside. At the completion of all testing, the used sand is autoclaved before reuse in future trials.

### Foraging Arena Construction

The foraging arena, tunnel and testing cage were custom built with acrylic plastic (Delvies Plastics, Salt Lake City, UT, USA). The 14 cm long tunnel enters the platform from underneath through one of the five 5.5 cm diameter holes in the arena platform. The platform is 8 cm above stage level and is made from 0.5 cm thick white Plexiglas. The arena is made from a transparent Plexiglas tube and the 0.4 cm thick walls raise 42 cm above the platform. The arena has a diameter of 35 cm. The walls of the arena were roughened with sand paper to limit glaring and recording artifacts.

### Arena Tracking Zones

The arena is organized into zones that are used to breakdown the behavior and foraging strategies used by each animal in the assay as detailed in Figure S1. Here, we provide additional details regarding the definition of these zones. The arena is divided into five sectors and the boundary of each sector is the mid point between two pots (see Figure S1). The arena is further divided into three concentric circles, including the middle center zone, the intermediate zone and the outer wall zone. The outer radius of the *Intermediate* zone intersects with the center of the pots and tunnel entry. The radius of the *Center* zone is half the radius of the *Intermediate* zone. A zone is also created around each pot in the arena. *Pot* zones have a radius of 1.7x the radius of the pot itself. Finally, to learn about the behavior of the animal related to entries to and exits from the arena, we define zones around the tunnel entry. The *Tunnel Entry* zone aligns with the entry hole of the actual tunnel. The *In Tunnel* zone is covering the most peripheral area of the *Tunnel Entry* and tracks the mouse just before leaving the arena completely. Whenever the mouse is in the cage, the tracking system is recording the mouse as being in the *In Tunnel* zone. The *Tunnel Zone* area has the same radius as the *Pot* zones.

### Automated Tracking

At the start of the trial, the tracking begins with a 10 second delay to allow time for the connection of the testing-cage to the arena. The mouse is first tracked when it appears in the *In Tunnel zone* (Figure 1) and position and movement is continuously recorded after this time point until the end of the 30 min Exploration or Foraging Phase. The XY position of the center of the mouse is video tracked with at a rate of 30 frames per second. For the data analysis, tracking data from 0-25 minutes are used. Time spent in each zone, latency to visit a zone and number of visits to each zone, as well as the distance traveled, are calculated using the Ethovision software. The data is exported as results for the total 25-minute trial duration, as well as in 5 minute time bins. Sand displacement and food consummation measures are collected and calculated manually.

### Behavioral Measures

All 152 measures captured during Exploration or Foraging phases of the assay are presented in Table S1. Each measure is assigned a custom ID, as indicated in the table. To calculate the time spent in the *Tunnel Entry* zone, time in the *In Tunnel zone* is subtracted from the raw time spent in the *Tunnel Entry zone*. To calculate the time spent in *Tunnel Zone*, the raw time in the Tunnel Entry zone is subtracted from the raw time spent in *Tunnel* z*one*. To calculate the total time spent in the arena, the time spent in the *In Tunnel* zone is subtracted the raw time in the arena. For calculation of these measures for specific time bins, the corresponding measures for each time 5 min time bin output by Noldus Ethovision are used instead of the total time measures. For the time spent in all other zones in the arena, the Noldus Ethovision output was used directly for both total time and time binned results. The latency to appear on the platform is scored manually, by selecting the last video frame when the center of the mouse is tracked in the *Tunnel Entry* zone before climbing onto the arena and standing with all four paws on the platform.

### Behavioral Measure Data Normalization

As indicated in Table S1, all ‘time spent in zone’, sand displaced, food consumed and zone visit measures were normalized to the total time spent in the arena (TTA) by dividing each value by TTA (x/TTA). For time bin values, the TTA for the corresponding time bin is used for normalization (i.e. x-1/TTA-1). All latencies to visit pots were normalized to the latency to enter the platform (LEP) by subtracting the LEP (x-LEP). Latency to the center zone after arena entry (LCAE) is already normalized to LEP and does not need any further normalization. All percentage measures do not need any normalization. Time on the platform itself and time in the tunnel (cage) are not included in the normalized dataset because they are closely related and redundant to TTA.

### Locomotor Measures

The raw data files generated by Noldus Ethovision listing all XY coordinated and all zones as well as distance traveled and velocity for each frame were used to extract data describing excursions and locomotor patterns using custom code, including the duration and number of bouts at different velocities. An excursion is defined as beginning when the mouse leaves the tunnel (In Tunnel zone) and ending when the mouse returns to the tunnel (one round trip). The minimum number of frames for an excursion sequence is 10. Continuous velocity values in the data were categorized into three velocity classes, *slow*: velocity <= 5 equals; *medium*: velocity >5 <= 15; *fast:* velocity >15. The length of the bout is calculated using the number of frames in the sequence and all sequences of the same velocity class longer than 3 frames are counted as a single velocity bout.

### DeepFeat and Data Analysis

**See Detailed Protocol Folder**

